# The evolution of ecological flexibility, large brains, and long lives: capuchin monkey genomics revealed with fecalFACS

**DOI:** 10.1101/366112

**Authors:** Joseph D. Orkin, Michael J. Montague, Daniela Tejada-Martinez, Marc de Manuel, Javier del Campo, Saul Cheves Hernandez, Anthony Di Fiore, Claudia Fontsere, Jason A. Hodgson, Mareike C. Janiak, Lukas F.K. Kuderna, Esther Lizano, Maria Pia Martin, Yoshihito Niimura, George H. Perry, Carmen Soto Valverde, Jia Tang, Wesley C. Warren, João Pedro de Magalhães, Shoji Kawamura, Tomàs Marquès-Bonet, Roman Krawetz, Amanda D. Melin

**Author notes:** Corresponding authors Joseph Orkin, UPF-PRBB, Carrer Dr. Aiguader, 88, Barcelona, 08003, Spain, Amanda Melin, University of Calgary, 2500 University Dr. NW. T2N 1N4, Calgary, AB, Canada.

## Abstract

Ecological flexibility, extended lifespans, and large brains, have long intrigued evolutionary biologists, and comparative genomics offers an efficient and effective tool for generating new insights into the evolution of such traits. Studies of capuchin monkeys are particularly well situated to shed light on the selective pressures and genetic underpinnings of local adaptation to diverse habitats, longevity, and brain development. Distributed widely across Central and South America, they are inventive and extractive foragers, known for their sensorimotor intelligence. Capuchins have the largest relative brain size of any monkey and a lifespan that exceeds 50 years, despite their small (3-5 kg) body size. We assemble a *de novo* reference genome for *Cebus imitator* and provide the first genome annotation of a capuchin monkey. Through high-depth sequencing of DNA derived from blood, various tissues and feces via fluorescence activated cell sorting (fecalFACS) to isolate monkey epithelial cells, we compared genomes of capuchin populations from tropical dry forests and lowland rainforests and identified population divergence in genes involved in water balance, kidney function, and metabolism. Through a comparative genomics approach spanning a wide diversity of mammals, we identified genes under positive selection associated with longevity and brain development. Additionally, we provide a technological advancement in the use of non-invasive genomics for studies of free-ranging mammals. Our intra- and interspecific comparative study of capuchin genomics provides new insights into processes underlying local adaptation to diverse and physiologically challenging environments, as well as the molecular basis of brain evolution and longevity.

**SIGNIFICANCE:** Surviving challenging environments, living long lives, and engaging in complex cognitive processes are hallmark characteristics of human evolution. Similar traits have evolved in parallel in capuchin monkeys, but their genetic underpinnings remain unexplored. We developed and annotated a reference assembly for white-faced capuchin monkeys to explore the evolution of these phenotypes. By comparing populations of capuchins inhabiting rainforest versus dry forests with seasonal droughts, we detected selection in genes associated with kidney function, muscular wasting, and metabolism, suggesting adaptation to periodic resource scarcity. When comparing capuchins to other mammals, we identified evidence of selection in multiple genes implicated in longevity and brain development. Our research was facilitated by our new method to generate high- and low-coverage genomes from non-invasive biomaterials.

## BACKGROUND

Large brains, long lifespans, extended juvenescence, tool use, and problem solving are hallmark characteristics of great apes, and are of enduring interest in studies of human evolution (1–4). Similar suites of traits have arisen in other lineages, including some cetaceans, corvids and, independently, in another radiation of primates, the capuchin monkeys. Like great apes, they have diverse diets, consume and seek out high-energy resources, engage in complex extractive foraging techniques (5, 6) to consume difficult-to-access invertebrates and nuts (6), and have an extended lifespan, presently recorded up to 54 years in captivity (7, 8). While they do not show evidence of some traits linked with large brain size in humans, e.g. human-like social networks and cultural and technological transmission from older to younger groupmates, their propensity for tool use and their ecological flexibility may have contributed to their convergence with the great apes (9) offering opportunities for understanding the evolution of key traits via the comparative method (10–12). Similar approaches have revealed positive selection on genes related to brain size and long lives in great apes and other mammals (13, 14), but our understanding of the genetic underpinnings remains far from complete.

Capuchins also offer excellent opportunities to study local adaptation to challenging seasonal biomes. They occupy diverse habitats, including rainforests and, in the northern extent of their range, tropical dry forests. Particular challenges of the tropical dry forest are staying hydrated during the seasonally prominent droughts, high temperatures in the absence of foliage, and coping metabolically with periods of fruit dearth (Figure 1). The long term study of white-faced capuchins (*Cebus imitator*) occupying these seasonal forests has demonstrated that high infant mortality rates accompany periods of intense drought, illustrating the strength of this selective pressure (15). Furthermore, the seasonally low abundance of fruit is associated with muscular wasting and low circulating levels of urinary creatinine among these capuchins (16). Additionally, the sensory challenges of food search in dry versus humid biomes are also distinct. Odor detection and propagation is affected by temperature and humidity (17), and color vision is hypothesized to be adaptive in the search for ripe fruits and young reddish leaves against a background of thick, mature foliage (18), which is absent for long stretches in dry deciduous forests. The behavioral plasticity of capuchins is widely acknowledged as a source of their ability to adapt to these dramatically different habitats (19–21). However, physiological processes, including water balance and metabolic adaptations to low caloric intake, and sensory adaptations to food search, are also anticipated to be targets of natural selection, as seen in other mammals (22–24). Understanding population-level differences between primates inhabiting different biomes, contextualized by their demographic history, genomic diversity, and historical patterns of migration, will generate new insights.

**Figure 1:**
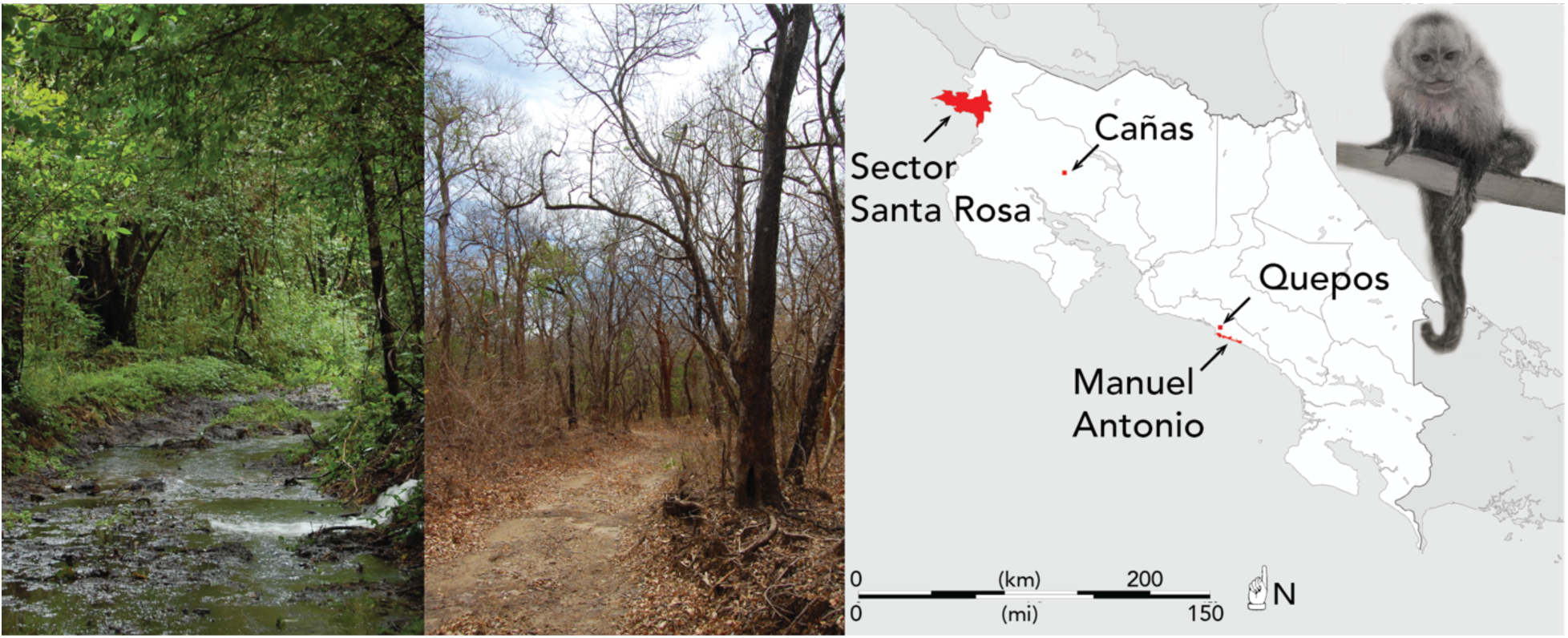
Sector Santa Rosa (SSR) during wet (left) and dry (middle) seasons. Right: Map of sampling locations in Costa Rica. The two northern sites, SSR and Cañas, have tropical dry forest biomes, whereas the two southern sites, Quepos and Manuel Antonio are tropical wet forests. Photos - Amanda Melin; Drawing of white faced capuchin monkey - Alejandra Tejada-Martinez; Map: Eric Gaba – Wikimedia Commons user: Sting

Unfortunately, high quality biological specimens from wild capuchins are not readily available. As is the case with most of the world’s primates, many of which are rare or threatened (25), this has limited the scope of questions about their biology that can be answered. Although recent advances in non-invasive genomics have allowed for the sequencing of partial genomes by enriching the proportion of endogenous DNA in feces (26–29), it has not yet been feasible to sequence whole genomes from non-invasive samples at high-coverage; this has limited the extent to which non-invasive samples can be used to generate genomic resources for non-model organisms, such as capuchins.

Toward identifying the genetic underpinnings of local adaptation to seasonally harsh environments, large brains, and long lifespans, we assembled and annotated the first reference genome of *C. imitator* (SI Appendix: Table S1). Additionally, we sequenced the genomes of individuals inhabiting two distinct environments in Costa Rica: lowland evergreen rainforest (southern population) and lowland tropical dry forest (northern population). We conducted high-coverage re-sequencing (10X - 47X) for 10 of these individuals, and sequenced an additional 13 at low-coverage (0.1X - 4.4X). Importantly, to facilitate the population-wide analyses without the need for potentially harmful invasive sampling of wild primates, we developed a novel method for minimally-biased, whole-genome sequencing of fecal DNA using fluorescence-activated cell sorting (fecalFACS) that we used to develop both high and low-coverage genomes (Figure 2). With these genomes, we assess the genetic underpinnings of capuchin-specific biology and adaptation in a comparative framework. First, we scanned the high-coverage genomes (6 from the northern dry forest, and 4 from the southern rainforest) for regions exhibiting population specific divergence to assess the extent of local adaptation to dry forest and rainforest environments. We examine how genes related to water balance, metabolism, muscular wasting, and chemosensation have diverged between populations. Second, we conduct an analysis of positive selection on the white-faced capuchin genome through codon-based models of evolution and enrichment tests focusing on genes that may underlie brain development and lifespan. Third, we identify the population structure, genomic diversity, and demographic history of the species using a mixture of traditional and non-invasive fecalFACS genomes (n=23).

**Figure 2:**
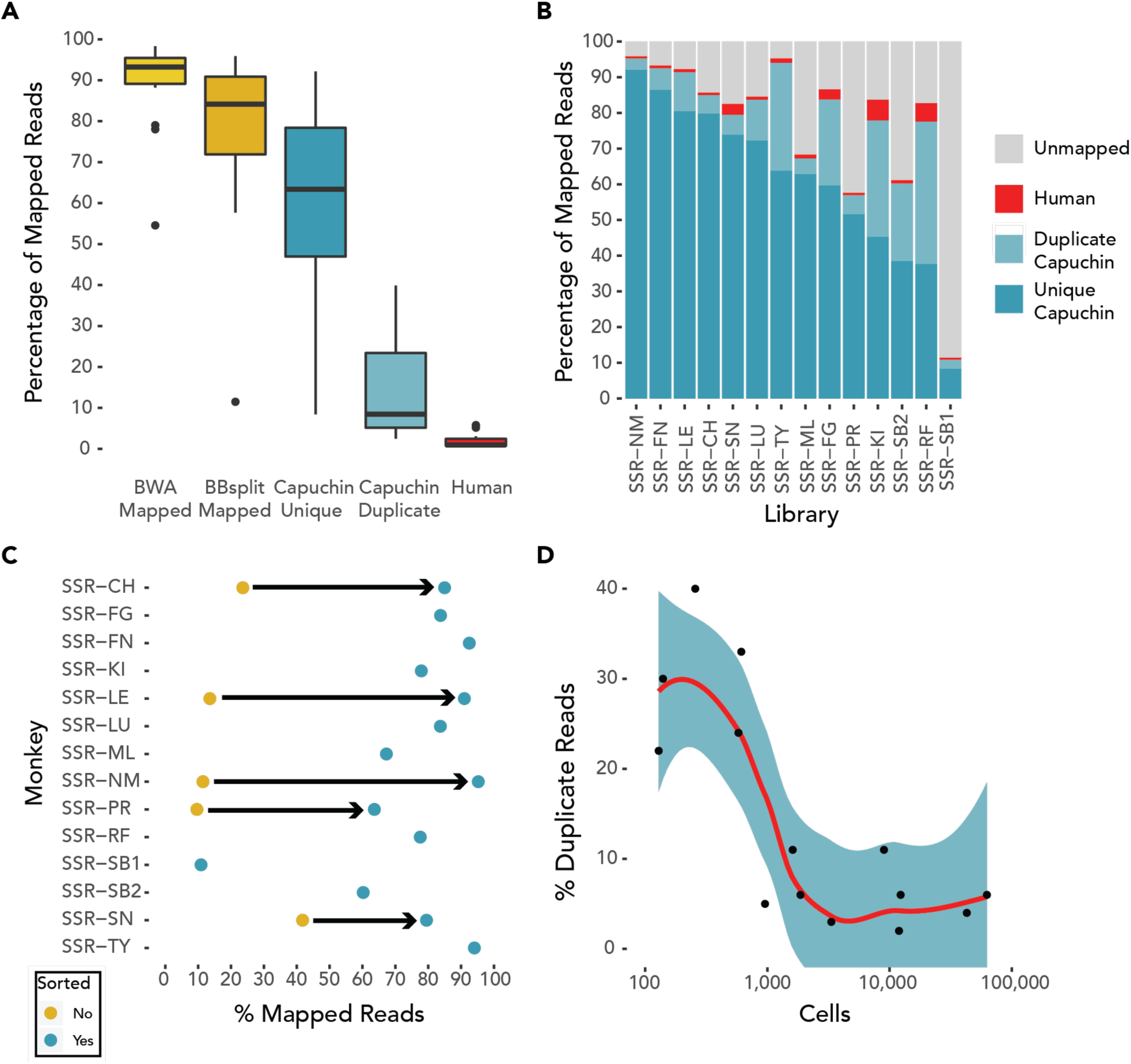
Mapping percentages of sequencing reads from RNAlater preserved fDNA libraries prepared with FACS for A) all samples [Box-plot elements: center line, median; box limits, upper and lower quartiles; whiskers, 1.5x interquartile range; points, outliers], and B) individual libraries. C) Increase in mapping rate for RNAlater preserved samples. D) Relationship between mapped read duplication and number of cells with LOESS smoothing. The duplicate rate decreases sharply once a threshold of about 1,000 cells is reached.

## RESULTS

### Local adaptation to seasonal food and water scarcity

We predicted that genes related to water balance, metabolism, muscular wasting, and chemosensation would differ between dry forest and rainforest populations of white-faced capuchins, reflecting local adaptation to different habitats. To test this, we searched for associations between genes in windows with high F_ST_ and divergent non-synonymous single nucleotide variants (SNV)s with high or moderate effect. Of the 299 genes identified in high F_ST_ windows, 39 had highly differentiated (F_ST_ >= 0.75) non-synonymous SNVs (SI Appendix: Figure S1, Table S2; SI Dataset). We identified 26 genes with non-synonymous SNVs of high or moderate effect that are fixed between populations in our dataset; 13 of these overlapped with the set of 39. Our enrichment analysis identified a single significant GO biological process: regulation of protein localization to cilium (GO: 1903564). In accordance with our hypothesis, disruptions of cilia proteins are predominantly associated with disorders of the kidney and retina (ciliopathies) (30). Furthermore, enrichment of genes associated with disease states was linked to kidney function, metabolism, and muscular wasting. These genes are good candidates for adaptive resilience to seasonal water and food shortages and warrant further investigation. We highlight several genes of particular promise (Figure 3, SI Appendix: Figure S2).

**Figure 3:**
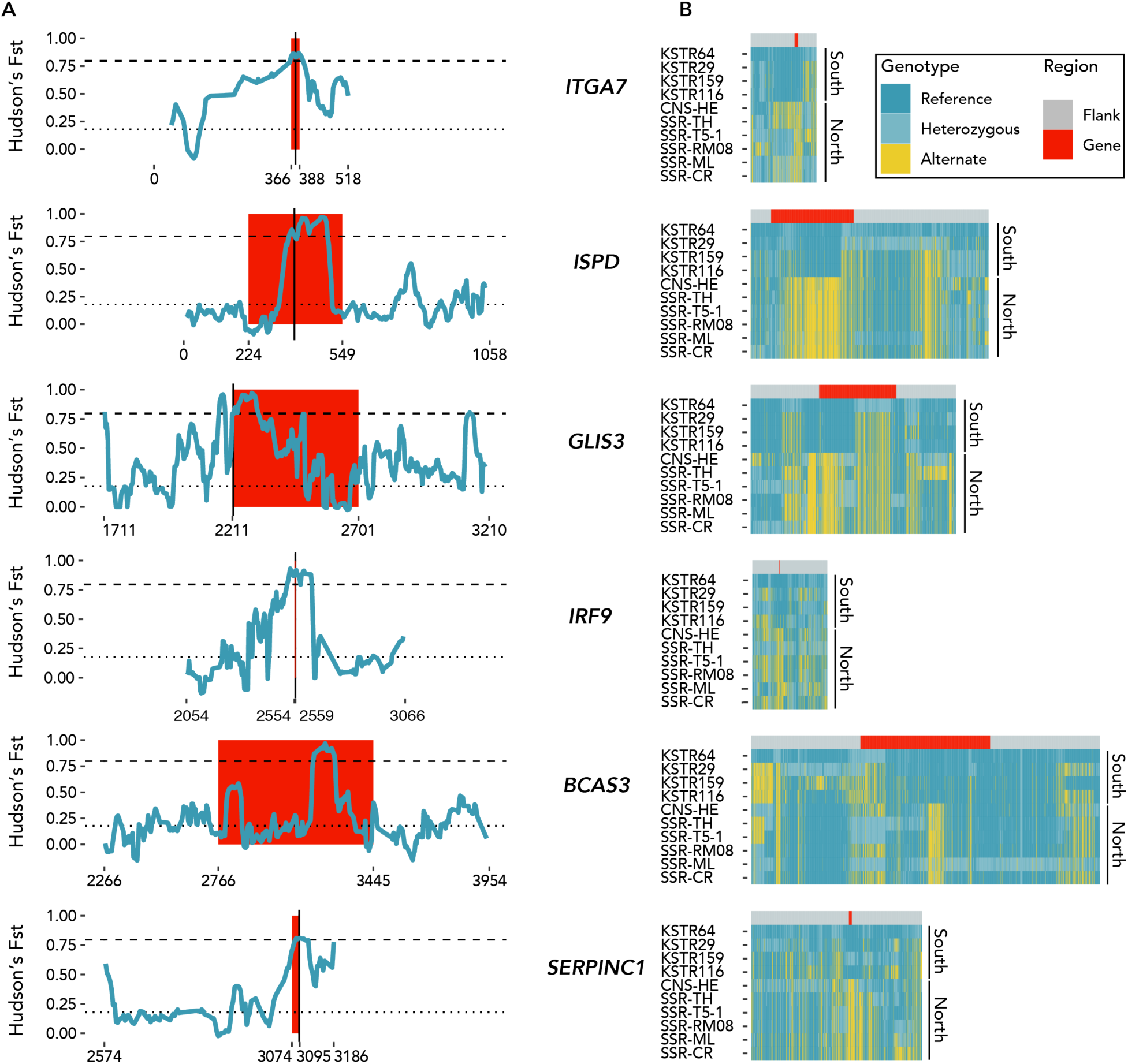
Highly differentiated genes between wet and dry forest populations involved in diabetes, kidney function, and creatinine levels. A: Hudson’s F_ST_ within windows of 20kb with a 4 kb slide. Gene regions are in red, flanked by 500 kb (or length to beginning or end of scaffold) of sequence. X-axis values correspond to position along the scaffold. The dotted line indicates average F_ST_ value across all windows (F_ST_ = 0.178), and the dashed line represents the top 0.5% of values (F_ST_ = 0.797). Vertical black lines indicate a non-synonymous SNP with an Fst >= 0.750, excluding BCAS3 (see Results). B: Heatmaps indicating the pattern of SNP variation within and surrounding highly divergent genes. SNVs within the genes are located under the red band and those within 200 kb of flanking region under the gray bands.

#### Evidence of adaptation to food and water scarcity

Population differences in multiple candidate genes indicate that dry-forest capuchins could be adapted to seasonal drought-like conditions and food scarcity. *SERPINC1* encodes antithrombin III, which is involved in anticoagulant and anti-inflammatory responses associated with numerous kidney-related disorders including salt-sensitive hypertension, proteinuria, and nephrotic syndrome (31–34). Additionally, sequence variants of *AXDND1* (as identified in the GeneCards database) are associated with nephrotic syndrome. *BCAS3* is expressed in multiple distal nephron cells types (35), and is associated with four pleiotropic kidney functions (concentrations of serum creatinine, blood urea nitrogen, uric acid, and the estimated glomerular filtration rate based on serum creatinine level) (36). Although we did not identify any population-specific non-synonymous SNVs in *BCAS3*, the genomic windows encompassing the gene rank among the highest regions of F_ST_ in our dataset, and intronic variation in BCAS3 putatively impacts estimated glomerular filtration rate in humans (35).

We also examined candidate genes associated with metabolism and with catalysis of muscle tissues, which occurs when energy intake is insufficient to fuel metabolic and energetic needs over long periods. We found population differentiation in genes associated with congenital muscular dystrophies--including mild forms that present with muscular wasting--and abnormal circulating creatine kinase concentration (HP:0040081): *ITGA7, ISPD* (*CRPPA*), and *SYNE2* (37–40). The gene *LAMA2* (41), which falls in a high F_ST_ window, has also been associated with muscular pathologies, some of which can be reduced by transgenic overexpression of *ITGA7*. Finally, we examined genes with roles in carbohydrate and lipid metabolism. We observed differentiation in *GLIS3*, which is associated with type 1, type 2, and neonatal diabetes (42). IRF9, which also has a fixed SNV of moderate effect between populations, has been shown to influence glucose metabolism, insulin sensitivity, and obesity, in knockout mice (43). *IRF9* has also been associated with immunity and susceptibility to viral infections (OMIM: 618648).

Given the appearance of both diabetes and kidney disorders in our gene sets, we conducted an *a posteriori* search of our high F_ST_ gene set for overrepresentation of genes associated with diabetic nephropathy (EFO_0000401) in the GWAS catalogue (SI Appendix: Table S3). Seven genes were present in both our gene set of 299 high F_ST_ genes and the diabetic nephropathy set of 117. Given the 16,553 annotated genes with HGNC Gene IDs, seven overlapping genes would occur with p = 0.00046 when permuted 100,000 times.

#### Evidence of adaptation in sensory systems

Intact gene sequences were found for two opsin genes, *OPN1SW* and *OPN1LW*, which underlie color vision and acuity. In both populations, we observed similar levels of polymorphism in the long-wavelength sensitive opsin gene (*OPN1LW*), with the same allelic variants at each of the three long wave cone opsin tuning sites (180, A/S; 277, F/Y; and 285, T/A) (SI Appendix: Table S4). We observed some evidence for population specific variation associated with the photoreceptive layers of the retina; a fixed non-synonymous SNV in *CCDC66*, which falls in a high F_ST_ region, is highly expressed in photoreceptive layers of the retina (44). Turning to olfaction, we identified 614 olfactory receptor (OR) genes and pseudogenes in the capuchin reference genome: 408 intact, 45 truncated, and 161 pseudogenized (SI Appendix: Table S5). To test for population differences in the OR gene repertoire, we assembled each olfactory gene/pseudogene independently in each individual. The proportion of total functional ORs was stable across individuals and populations, with seemingly trivial fluctuations (North 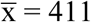, s = 1.6; South 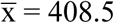, s = 1.3), possibly driven by a small difference in OR family 5/8/9 (SI Appendix: Table S6). We also identified 7 intact vomeronasal (VR) and 28 taste receptor (TASR) and taste receptor-like genes (SI Appendix: Table S7), two of which (*TAS1R* and *TAS2R4*) have non-synonymous SNVs with fixed variants in the north (SI Appendix: Table S8).

### Evolution of brain development and longevity in capuchins

To identify genes that underlie brain development and lifespan, which are of particular interest given the derived features of capuchin biology, we ran a codon-based positive selection analysis with PAML and subsequently tested for functional enrichment with ToppFunn (45) (Figure 4). Of the genes identified as being under positive selection in the branch-site model (748 genes), and the branch model (612 genes), we identified significant enrichment in genes associated with brain development (52 genes: SI Dataset), neurogenesis (77 genes: SI Dataset), and microcephaly (31 genes: SI Dataset). For example, *WDR62, BPTF, BBS7, NUP113*, mutations are directly associated with brain size and related malformations, including microcephaly (46–49). *MTOR* signaling malfunction is also implicated in developmental brain malformations (50), and *NUP113* is involved in nuclear migration during mammalian brain development (51). Several genes are linked with brain tumor formation (including *ZNF217*), and others with cognitive ability (e.g. *PHF8 (52)*).

**Figure 4:**
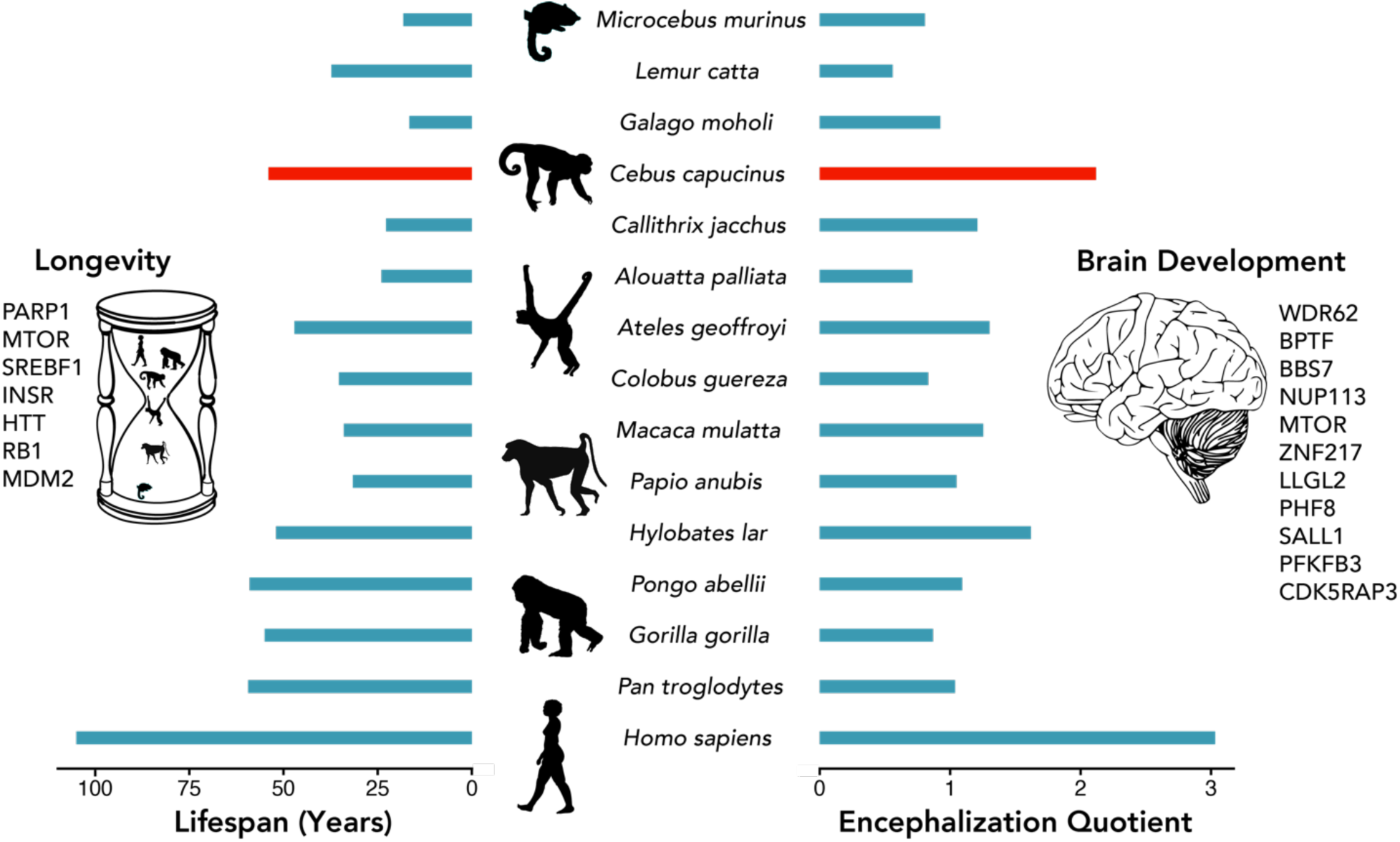
Genes under positive selection in white-faced capuchin monkeys that are associated with maximum recorded longevity in captivity and brain development. Values from *Cebus capucinus* are used in place of *Cebus imitator*, given the recent taxonomic split. We calculated encephalization quotient as brain mass/(0.085*(body mass^0.775^)) (1), with data from (1). Lifespan values are from (8).

We found 48 genes putatively linked to longevity, as identified in the GenAge and CellAge databases (14, 53), to be under positive selection in capuchins including *PARP1, MTOR, SREBF1, INSR, HTT, RB1* and *MDM2* (SI Appendix: Table S9). Of note, poly (ADP-ribose) polymerase 1 (PARP1) putatively serves as a determinant of mammalian aging due to its activity in the recovery of cells from DNA damage, and *MTOR* acts as a regulator of cell growth and proliferation while also being generally involved in metabolism and other processes. Additional key genes in aging and metabolism include sterol regulatory element binding transcription factor 1 (*SREBF1*), which acts as a regulator of the metabolic benefits of caloric restriction (54, 55), and the insulin receptor (*INSR*), a major player in longevity regulation (56). As for specific age-related diseases, huntingtin (*HTT*) is under selection in mammals; *HTT* is not only involved in Huntington’s disease but has also been associated with longevity in mice (57). Lastly, various cell cycle regulators (e.g., *RB1, MDM2*) are also under positive selection in capuchins, and indeed, cell cycle is an enriched term among positively selected genes (SI Appendix: Table S9), though these could be related to other life history traits like developmental schedules that correlate with longevity.

### Population genomics of Costa Rican capuchins with fecalFACS

The pattern of clustering in our maximum likelihood SNV tree recapitulates the expected patterns of geographic distance and ecological separation in our samples (Figure 5). Likewise, in the projected PCA all individuals from the seasonal dry forests in the northwest are sharply discriminated from individuals inhabiting the southern rainforests along PC1. Levels of heterozygosity calculated in overlapping 1 Mb genomic windows (with a step size of 100 kb) were significantly higher in the southern population (W = 1,535,400,000, p-value < 2.2e-16; Figure 6A, SI Appendix: Figure S3). Furthermore, the median pairwise heterozygosity for each southern individual (range: 0.00065 - 0.00071) was higher than any northern monkey (0.00047 - 0.00057) (W = 0, p-value = 0.009524; SI Appendix: Table S10). In the northern population, we also identified long runs of homozygosity significantly more often (W = 24, p-value = 0.009524), and more of the longest runs (>= 5 Mb) (W = 1315.5, p-value = 0.03053; Figures 6B,; SI Appendix: Figures S4, S5). Pairwise sequential Markovian coalescent (PSMC) analysis of demographic history (Figure 6C) reveals that white-faced capuchins had a peak effective population size of ∼60,000 effective individuals ∼1 Ma, which declined to fewer than 20,000 during the middle to late Pleistocene transition. After recovering during the middle Pleistocene, they declined precipitously through the late Pleistocene to fewer than 5,000 effective individuals.

**Figure 5:**
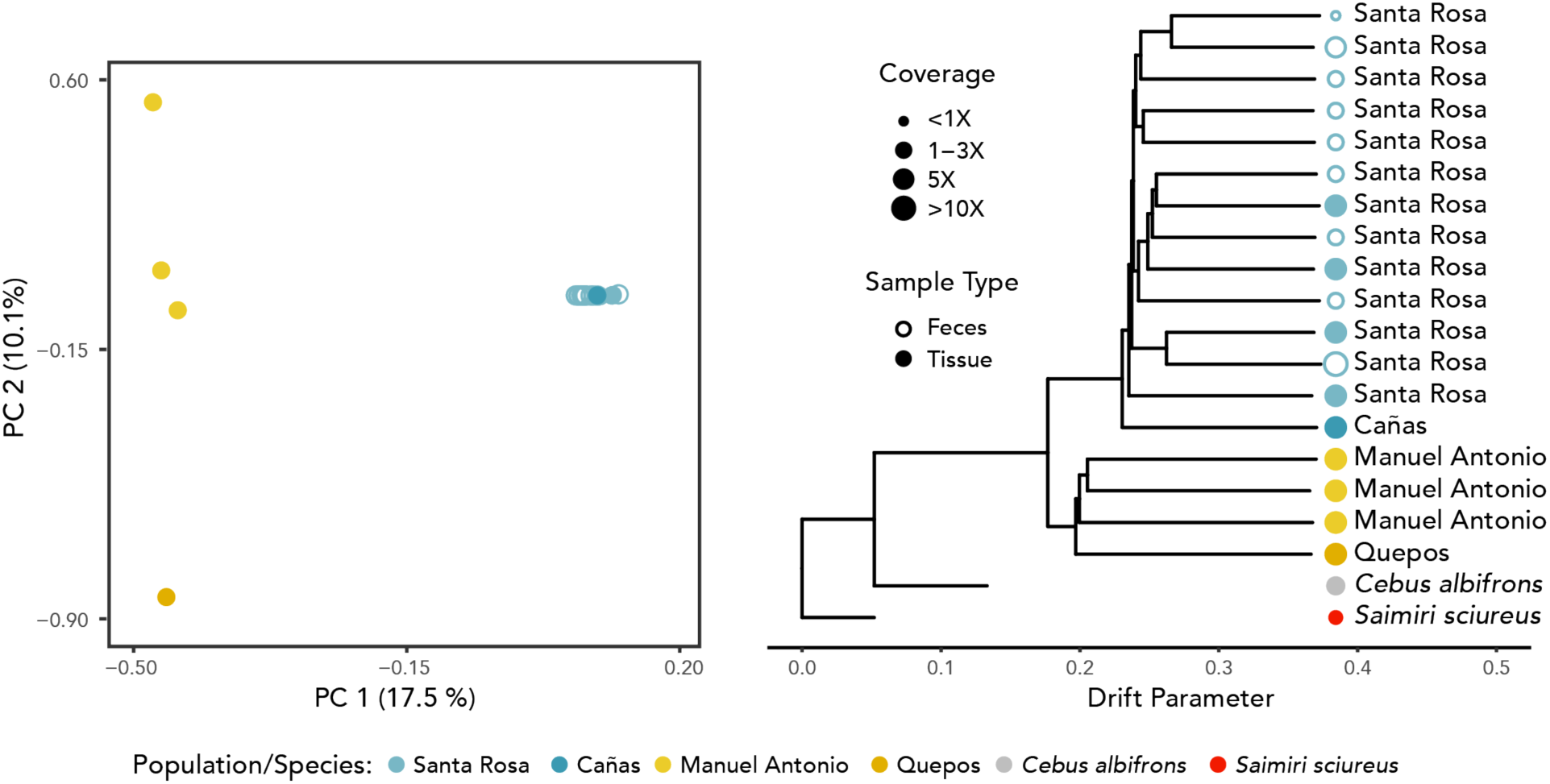
Population subdivision in *Cebus imitator*. Left: Principal components of 13 fecal and 10 blood/tissue libraries from white faced capuchins. Individuals from northern and southern sites separate on PC 1. Low- and high-coverage *C. imitator* samples from Santa Rosa plot in the same cluster. Right: Maximum likelihood tree of 9 fecal and 10 blood/tissue libraries from *C. imitator* (samples with less than 0.5X coverage were excluded). Among the white-faced capuchin samples, individuals from northern (dry forest) and southern (wet forest) regions form the primary split; secondary splits reflect the individuals from different sites within regions. The short branch lengths of the outgroups are a result of only polymorphic positions within *C. imitator* being used to construct the tree.

**Figure 6:**
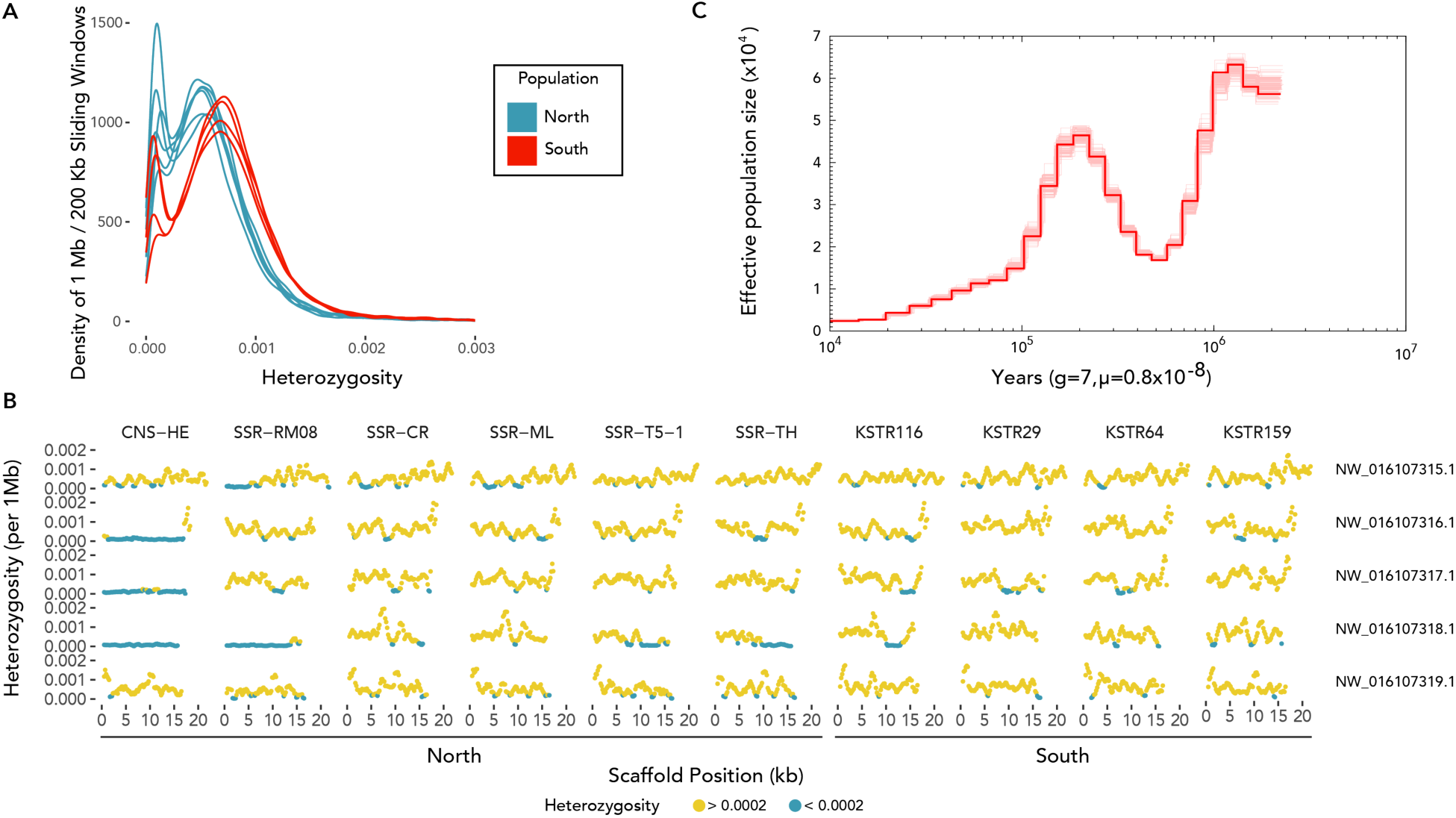
A: Density plot of 1 Mb windows with a slide of 200 kb in northern and southern populations. The distribution of windows from the northern population indicates lower heterozygosity than the southern distribution. The individuals from the southern population show consistently higher values. B: Long runs of homozygosity in the 5 largest scaffolds. Blue dots represent windows with depleted heterozygosity. The individuals with the longest runs of homozygosity come from the northern population. C: PSMC plot of effective population size over time.

Of the 25 capuchin DNA samples (comprising 23 individuals) that we sequenced and mapped to our reference genome, 16 were fecal-derived (SI Appendix: Table S11). This includes the first--to our knowledge--high-coverage (12.2 X) whole mammalian genome generated from a fecal sample. When comparing the high-coverage tissue-derived genome from the Santa Rosa site to those generated from our novel application of fluorescence-activated cell sorting to isolate fecal-sourced cells (fecalFACS), we observed no substantial difference in quality, coverage, heterozygosity, or GC content (SI Appendix: Figures S6, S7). FecalFACS yielded high mapping (median 93%) and low duplication rates (median 6%) (Figure 2; details in Methods). Population clustering relationships were not perturbed by depth of coverage, or source material (tissue-based vs fecalFACS genomic libraries) (Figure 5, SI Appendix: Table S12).

## DISCUSSION

### Local adaptation to seasonal environments

Patterns of genetic differentiation are consistent with our hypothesis that capuchins inhabiting the tropical dry forests have undergone local adaptation to the extreme seasonality of rainfall and food availability. We observed population-specific variation in genes implicated in water/salt balance, kidney function, muscle wasting, and metabolism. *SERPINC1* may reflect selection for drought resistance in the dry-forest capuchins. It has previously been linked to water/salinity balance in mammals; for example, it is overexpressed in the renal cortex of Dahl salt-sensitive rats fed a high salt diet (34). Additionally, the associations of *BCAS3* with glomerular filtration rate and *AXDND1* with nephrotic syndrome, a kidney disorder that commonly presents with edema and proteinuria, are consistent with this hypothesis. Regarding metabolism, *GLIS3* is the only gene other than *INS* (insulin) known to be associated with all three forms of diabetes (42), and *IRF9* has been shown to have a strong effect on body weight in mice, likely through its key role in glucose and lipid metabolism (43). Cautiously, we also point to *ABCC5*, which nearly missed our F_ST_ threshold, but does contain a fixed non-synonymous SNV in our two populations. The expression of *ABCC5* “plays a central role in energy metabolism in mammals,” affecting fat mass and insulin sensitivity in knockout mice and the onset of diabetes in humans (58). Additionally, the significant overabundance of genes associated with diabetic nephropathy in high F_ST_ windows is consistent with there being different functional pressures on kidney and metabolic function in these two populations.

That multiple genes in high F_ST_ are associated with muscle wasting and/or serum creatinine and creatinine kinase levels (*ITGA7, ISPD, SYNE2*, and *BCAS3*) is of particular interest. Creatinine is a byproduct of the metabolism of creatine phosphate in skeletal muscle, which is normally filtered by the kidneys. It has been used as a clinical biomarker of kidney function, chronic kidney disease (36), and as a monitor of muscle mass (59). Previous field study of urinary creatinine levels of wild capuchins from the northern population of Sector Santa Rosa (SSR) document expected relationships with decreased muscle mass during periods of seasonally low fruit availability, and highlight the biological impacts of seasonal food shortages in the dry forest (16). Study of seasonal variation in SSR capuchin gut microbes (60, 61) also provides evidence of periodic episodes of resource stress and poorer health that differs markedly from those of capuchins inhabiting nonseasonal forests (62). Finally, a recent study has documented mass infant mortality in SSR capuchins following years with longer-than average droughts (15). Together, these observations fit with the notion that animals living in seasonal environments, or pursuing seasonal migrations, are more likely to have weight fluctuations through binge-subsist cycles that map onto food abundance (63). We take this as promising evidence that these genes have been under selection in the northwestern population.

Given that selection operates on both gene function and regulation, we suspect the observed variation is affecting gene expression or enzymatic efficiency, which offers a promising avenue for future research. In addition, overlap between metabolic and immune functions in *IRF9* could also suggest inflammation as another target of selection. Because challenging environments will exert selective pressures on numerous other health-related bodily functions, including immunity, future study in this area may be instructive (64). In this vein, further research could examine *TLR4*, which has been proposed as a “molecular link among nutrition, lipids and inflammation,” because of its involvement not only with cytokine expression, but also glucose uptake, body mass, and insulin metabolism (65). Although *TLR4* fell outside our high F_ST_ window threshold, we did identify a fixed non-synonymous SNV between populations.

Turning to sensory systems, we found an intact short wavelength (blue) sensitive opsin gene *OPN1SW*, and three alleles of the long wavelength sensitive opsin gene (*OPN1LW*) that are consistent with previous reports for *Cebus* (66–68). Like most other primates in the Americas, capuchins possess a polymorphic color vision system, characterized by extensive intraspecific genotypic and phenotypic variation (69, 70). This widespread, persistent variation is a fascinating example of balancing selection that has captivated biologists for decades (67, 70, 71). That we find no evidence of differences in the number opsin alleles, nor novel variants, between dry forest and rainforest populations suggests that these habitat differences are not selecting for variation in color vision intraspecifically. However, we did observe population-specific variation in *CCDC66* that could be associated with scotopic (rod-driven) photoreceptor response. Electroretinography of *CCDC66* -/- mice reveals a significant reduction in scotopic photoreceptor response (44), indicating a potential effect on vision in low-light conditions. Curiously, *CCDC66* may impact smell as well as vision. *CCDC66* -/- mice, also display neurodegeneration of the olfactory bulb, and have reduced odor discrimination performance of lemon smells (72). The ecological significance of this result, if any, is unclear at present but may warrant future attention. Turning to chemosensory receptor genes, we find the numbers of functional OR, VR, and TASRs are similar to, or slightly higher than, the number identified in other anthropoid primates (73–76). Interestingly, the VR gene repertoire of capuchins highlights the persistent role of the vomeronasal organ that is used in social communication of other mammals and is likely important to the social relationships of capuchins, which frequently engage in urine washing and “hand-sniffing” behaviors (77). The functional significance of the TASR variants we identified is unknown, but may be revealed via cellular expression systems in future research (78). The tuning of the chemosensory receptors to specific stimuli could vary between habitat types and may be elucidated by future work.

A limitation of our study is the relatively small number of sampled individuals from few sampling sites, and that deeper sampling would reduce the chance of spurious results. In order to minimize the possibility of such random hits, we held to strict requirements when identifying evidence of local adaptation. We required that a function directly related to one of our *a priori* hypotheses be identified as significantly associated by ToppFunn in the strict set of 39 genes that were in the top 0.5% of F_ST_ windows and contained a SNV with high or moderate effect (e.g. *ITGA7, ISPD, SERPINC1, IRF9, GLIS3, CCDC66*) before examining the secondary gene sets (299 in high Fst regions and 26 with fixed SNVs) for supplementary support (e.g. *BCAS3, ABCC5, TLR4*).

### Evolution of brain development and longevity in capuchins

Among primates, capuchin monkeys are known for their relatively large brains, derived cognition, and sensorimotor intelligence (2, 5, 6, 9). Accordingly, it is perhaps unsurprising to see positive selection in the *Cebus* lineage and evidence of shifts in gene function linked to brain function and development relative to other primates. In particular, positive selection in *WDR62, BPTF, BBS7, NUP133*, and *MTOR*, and *PHF8* is consistent with observations that the capuchin lineage has undergone adaptation linked to brain development. The association of several of these genes with size related brain malformations (46–49), such as microcephaly, suggests that they could be influencing the large relative brain size of capuchins. Furthermore, the evidence of selection in PHF8, which is associated with human cognition, aligns with the link between brain size and intelligence that has been observed in other primates (79). While we highlight here the putative functional roles of these genes, which are based on clinical studies and comparative genomics, we acknowledge that further examination of their function in the context of capuchin biology is warranted.

In the context of longevity, it is noteworthy that we observed genes under selection associated with DNA damage response, metabolism, cell cycle and insulin signaling (13). Of particular interest are: *PARP1, MTOR, SREBF1, INSR1*, and *HTT*. Damage to the DNA is thought to be a major contributor to aging (80). Previous studies have also shown that genes involved in DNA damage responses exhibit longevity-specific selection patterns in mammals (14). It is therefore intriguing that *PARP1*, a gene suggested to be a determinant of mammalian aging (81), is under selection in capuchins. Another large body of research has associated the mechanistic target of rapamycin (*MTOR*) with aging and longevity in various organisms (82), making it a prime candidate for therapeutic interventions in aging (83). Other genomes of long-lived mammals also revealed genes related to DNA repair and DNA damage responses under selection (84, 85). Intriguingly, short-lived species also exhibit genes under selection related to insulin receptors, raising the possibility that the same pathways associated with aging in model organisms are involved in the evolution of both short- and long lifespans (86), an idea supported by our results. Of course, because aging-related genes often play multiple roles, for example in growth and development, it is impossible to be sure whether selection in these genes is related to aging or to other life-history traits, like growth rates and developmental times, that in turn correlate with longevity (87). Therefore, although we should be cautious about the biological significance of our findings, it is tempting to speculate that, like in other species, changes to specific aging-related genes or pathways, could contribute to the longevity of capuchins. Additionally, it is difficult to identify the evolutionary time depth beyond the *Saimiri* / *Cebus* branch (Estimated Time: 16.07 Mya; CI: 15.16 - 17.43 mya (88)) based on the present study. Future analyses of other capuchin species, in both extant genera--*Cebus* and *Sapajus*--along with other Cebids and platyrrhines will better reveal more recent selection. Future research assessing structural variation across the genome could also be fruitful, particularly in cases such as the copy number of CD33 related Siglec genes, which have previously been linked to longevity in mammals (89).

Finally, it is worth noting that subtle changes in the duration of expression of regulatory genes could have profound effects on both lifespan and brain development. Evidence of the temporal importance of the onset of neurogenesis and effects on interspecific variation in neuron composition are present in comparative study of owl and capuchin monkeys, for example (90). These changes are not possible to detect with the orthologous gene positive selection scan approach that we take here; future research examining likely differences in neuroembryological parameters, along with variation in gene expression profiles across the lifespan, would complement and extend these contributions.

### Population genomics with fecalFACS

We observe a clear demarcation in population structure between the northern dry- and southern wet-forest populations in Costa Rica. Higher levels of heterozygosity in the south and lower levels in the northwest are in accordance with the current hypothesis that capuchins dispersed northwards across Costa Rica. White-faced capuchins are the most northerly distributed member of the Cebinae, having dispersed over the isthmus of Panama in a presumed speciation event with *C. capucinus* in South America ∼1.6 Ma (91, 92). After expanding during the early late Pleistocene, white-faced capuchins appear to have undergone a dramatic reduction in effective population size. This pattern predates the movement of humans into Central America, and could reflect a series of population collapses and expansions caused by glacial shifts and fluctuating forest cover availability during the Pleistocene. *C. imitator* in SSR are near the northernmost limits of their range, which extends as far north as Honduras, and may represent some of the least genetically diverse members of their species. Given the limitations of the available sampling sites, it is possible that the appearance of an ecological divide is actually evidence of isolation by distance; however, given that the single individual from Cañas clusters closely with the individuals from SSR, despite a geographic distance of more than 100 km, we suggest that isolation by distance does not completely explain the population differentiation.

Through a novel use of flow cytometry/FACS, we successfully generated the first high-coverage, minimally biased mammalian genome solely from feces, along with a low-coverage SNP dataset that is suitable for population assignment and clustering. We also used the high-coverage fecalFACS genome to call SNPs (along with traditional blood and tissue genomes), demonstrating for the first time that non-invasively collected fecal samples can successfully be used to generate primate genomic resources. The clustering patterns in our trees and PCA plots do not reveal any samples that deviate from their expected geographic or ecological origin. These relationships are robust to both the coverage levels (< 1X to > 50X) and biological origins (feces, tissue, and blood) of the samples. The tight geographic clustering of individuals within the SSR sampling locale provides reasonable evidence that there is no substantial effect from fecalFACS on population structure. One possible limitation of fecalFACS could be the co-isolation of cells from prey items in carnivorous/omnivorous mammals. On rare occasions, capuchins eat small mammalian prey, such as infant coatimundis and squirrels (93), and we did observe a small contribution (< 2% of reads in two samples) that could be from mammalian prey (SI Appendix: Figure S8A). However, we successfully removed these reads with BBsplit (94) mapping (SI Appendix: Figure S8B), and we did not notice any effect of contamination in our results (Figure 5). We suspect that any dietary contribution is low, as partially digested cells would be more likely to be degraded in a way that further distinguishes them in size and granularity (i.e. the forward and side scatter parameters used in our FACS gating) from epithelial cells sloughed from the lining of the colon. Given that fecalFACS is cost-effective and minimizes the biases that commonly occur in traditional bait-and-capture approaches to the enrichment of endogenous DNA from feces, and does not require costly impractical preservation of biomaterial in liquid nitrogen but rather uses room-temperature stable storage in RNAlater, this method offers great benefits to the field of mammalian conservation and population genomics. Our approach is promising for future studies of wild animals where invasive sampling is not feasible or ethical.

## SUMMARY

In our population level analysis of wet- and dry-forest capuchins, we observed both evidence of population structure between and local adaptation to these different habitats. In particular, we identified selection in genes related to food and water scarcity, as well as muscular wasting, all of which have been observed during seasonal extremes in the dry forest population. Interspecific analyses revealed evidence of selection in *C. imitator* on genes involved with brain development and longevity. These results are in accordance with the remarkably long life span, large brain, and high degree of sensorimotor intelligence that has been observed in capuchins. These genes are good candidates for further investigation of traits which have evolved in parallel in apes and other mammals. Importantly, our insights were made possible through the first annotated reference assembly of a capuchin monkey and a novel use of flow cytometry/FACS to isolate epithelial cells from mammalian feces for population genomics. FecalFACS allowed us to generate both the first high-coverage, minimally biased mammalian genome solely from feces, as well as low-coverage SNV datasets for population level analyses.

## METHODS

### Study populations and sample collection

Central American white-faced capuchins *(Cebus imitator)*, a member of the gracile radiation of capuchins (genus *Cebus*), were recently recognized as a species, distinct from *C. capucinus* in South America (92). *Cebus imitator* occupies a wide diversity of habitats, spanning lowland rainforests and cloud forests in Panama and throughout southern, eastern, and central Costa Rica, and tropical dry forests in northwestern Costa Rica and Nicaragua. The annual precipitation and elevation of rainforest versus dry forest biomes in their current range vary dramatically, leading to considerable variation in the resident flora and fauna (95, 96). We sampled individual Costa Rican capuchin monkeys from populations inhabiting two distinct habitats. 1) lowland rainforest around Quepos, Puntarenas Province; and 2) tropical dry forest at two sites in Guanacaste Province. In total, we collected samples from 23 capuchins, a list of which is provided in SI Appendix: Table S11.

We sampled capuchins inhabiting a lowland tropical rainforest biome by collaborating with *Kids Saving the Rainforest* (KSTR) in Quepos, Costa Rica. We acquired blood samples from 4 wild capuchins from nearby populations who were undergoing treatment at the facility (although we were unable to collect paired fecal samples). For one of these individuals, an adult male white-faced capuchin that was mortally wounded by a vehicle in Costa Rica, we additionally sampled tissues from several organs. DNA derived from the kidney was used for the reference genome assembly.

We collected 21 samples from 19 individuals in the northern tropical dry forest. 16 fecal samples and 4 tissue samples were from free-ranging white-faced capuchin monkeys (*Cebus imitator*) at in the Sector Santa Rosa (SSR), part of the Área de Conservación Guanacaste in northwestern Costa Rica, which is a 163,000 hectare tropical dry forest nature reserve (Figure 1). Behavioral research of free-ranging white-faced capuchins has been ongoing at SSR since the 1980’s which allows for the reliable identification of known individuals from facial features and bodily scars (97). The 16 fresh fecal samples were collected from 14 white-faced capuchin monkeys immediately following defecation (SI Appendix: Table S10). We placed 1 mL of feces into conical 15 mL tubes pre-filled with 5 mL of RNAlater. RNAlater preserved fecal samples were sent to the University of Calgary, where they were stored at room temperature for up to three years. To evaluate other preservation methods, we also collected two additional capuchin monkey fecal samples (SSR-FL and a section of SSR-ML), which we stored in 1X PBS buffer and then frozen in liquid nitrogen with a betaine cryopreservative (98). Given the logistical challenges of carrying liquid nitrogen to remote field sites, we prioritized evaluation of samples stored in RNAlater. We also collected tissue and blood samples opportunistically. During the course of our study, 4 individual capuchin monkeys died of natural causes at SSR, from whom we were able to collect tissue samples, which were stored in RNAlater. Additionally, we collected a blood sample from 1 northern dry-forest individual housed at KSTR that originated near the town of Cañas, which is ∼100 km southwest of SSR.

### Summary of genome-wide sequencing, genome assembly and gene annotation

We assembled a reference genome for *Cebus imitator* from DNA extracted from the kidney of a male Costa Rican individual (KSTR64) using a short read approach (Illumina HiSeq 2500) with ALLPATHS-LG. To improve the quality of gene annotation, we isolated total RNA from the whole blood of an adult male white-faced capuchin (ID: CNS-HE) permanently residing at the KSTR wildlife rehabilitation center. The capuchin genome assembly was annotated with the NCBI pipeline previously described here: (http://www.ncbi.nlm.nih.gov/books/NBK169439/).

Our reference genome assembly for *Cebus imitator* is composed of 7,742 scaffolds (including single contig scaffolds) with an N50 scaffold length of 5.2 Mb and an N50 contig length of 41 kb. The final ungapped assembly length is 2.6 Gb (GenBank accession: GCA_001604975.1). Our estimate of total interspersed repeats using WindowMasker (99) output is 45.8%. The numbers of annotated genes are 20,740 and 9,556 for protein-coding and non-coding genes, respectively (SI Appendix: Table S1). Measures of gene representation using the known human RefSeq set of 56,230 transcripts show an average of >94% coverage with a mean identity of 92.5%. Overall, our draft assembly metrics and gene representation are consistent with other non-human primate (NHP) short-read reference assemblies (100). See SI Appendix for further methodological detail.

### Summary of positive natural selection analysis through codon-based models of evolution and enrichment tests

The phylogenetic arrangement in this study included 14 species as outgroups to *C. imitator*: three Platyrrhini (*Callithrix jacchus, Aotus nancymaae, Saimiri* boliviensis), six Catarrhini (*Macaca mulatta, Rhinopithecus roxellana, Nomascus leucogenys, Pan troglodytes, Homo sapiens, Pongo abelii*), one Strepsirrhini (*Microcebus murinus*), one rodent (*Mus musculus*), and three Laurasiatheria (*Canis lupus familiaris, Bos taurus*, and *Sus scrofa*). We identified 7,519 OGs present among the 15 species. We identified 612 genes under positive selection (p<0.05 after FDR correction) in the *Cebus* lineage using the branch model of codeml in PAML. We also performed a branch-site test and identified a second set of 748 genes under positive selection in *Cebus* (SI Dataset). We performed functional annotation analysis using ToppFunn in the ToppGene Suite (45) with default parameters. To ascertain which ontology processes the genes with signals of positive selection were involved, we focused the enrichment analysis on two functional categories: GO Biological Processes (BP) and the DisGeNET BeFree disease database. Finally, the genes with positive selection signal were intersected with the GenAge and CellAge databases (build 19,307 genes) (53). See SI Appendix for further methodological detail.

### FecalFACS

Before isolating cells by fluorescence-activated cell sorting, fecal samples were prepared using a series of washes and filtration steps. Fecal samples were vortexed for 30 s and centrifuged for 30 s at 2,500 g. Then the supernatant was passed through a 70 μm filter into a 50 mL tube and washed with DPBS. After transferring the resultant filtrate to a 15 mL tube, it was centrifuged at 1,500 RPM for 5 minutes to pellet the cells. Then we twice washed the cells with 13 mL of DPBS. We added 500 μL of DPBS to the pellet and re-filtered through a 35 μm filter into a 5 mL FACS tube. We prepared a negative control (to control for auto-fluorescence) with 500 μL of DPBS and one drop of the cell solution. To the remaining solution, we added 1 μL of AE1/AE3 Pan Cytokeratin Alexa Fluor® 488 antibody (ThermoFisher: 53-9003-82) or TOTO-3 Iodide (642/660) DNA stain, (ThermoFisher T3604) which we allowed to incubate at 4°C for at least 30 minutes.

We isolated cells using a BD FACSAria™ Fusion (BD Biosciences) flow cytometer at the University of Calgary Flow Cytometry Core. To sterilize the cytometer’s fluidics before processing each sample, we ran a 3% bleach solution through the system for four minutes at maximum pressure. We assessed background fluorescence and cellular integrity by processing the negative control sample prior to all prepared fecal samples. For each sample we first gated our target population by forward and side scatter characteristics that were likely to minimize bacteria and cellular debris (SI Appendix: Figure S9). Secondary and tertiary gates were implemented to remove cellular agglomerations. Finally, we selected cells with antibody or DNA fluorescence greater than background levels. In cases when staining was not effective, we sorted solely on the first three gates. Cells were pelleted and frozen at -20°C.

We isolated a median of 2,206 cells, with a range of 129 - 62,201 (SI Appendix: Table S12). The total amount of DNA per sample was low, ranging from 2.96 to 21.50 ng, with a median value of 7.85 ng, but in each case allowed for the successful generation of a sequencing library (SI Appendix: Table S12). The number of cells was not significantly correlated with the amount of extracted DNA (R=0.227; 95% CI (−0.345, 0.676); t=0.808, p > 0.05) or mapping rate (R = -0.204; 95% CI (−0.663, 0.367); t = -0.721; p > 0.05). Median mapping rates reached 93% (range: 55 - 98%) with BWA-MEM and 82% (range: 11 - 95%) with the more stringent BBsplit settings (Figure 2, SI Appendix: Table S11). Read duplication levels were low, with a median value of 6% (range: 2 - 40%) resulting in 63% (range: 8 - 92%) of reads being unique and mapping to the *Cebus imitator* 1.0 genome. The amount of duplicate reads was distributed bimodally across individuals, with reads from five samples having substantially higher duplication rates than the remaining nine. The rate of duplication was significantly correlated (R = -0.751; 95% CI (−0.917, -0.366); t = -3.94; p < 0.01) with the number of cells (log10 transformed), decreasing sharply above a threshold of about 1,000 cells (Figure 2). By sorting fecal samples with FACS, we substantially increased the percentage of reads mapping to the target genome. We selected five samples at random (SSR-CH, SSR-NM, SSR-LE, SSR-PR, SSR-SN) to compare pre- and post-FACS mapping rates. The mapping rates of unsorted feces ranged from 10 - 42%, with a median of 14% (Figure 2C). After flow sorting aliquots of these fecal samples, we obtained significantly higher mapping rates (V = 15, p < 0.05) for each sample, ranging from 64 - 95%, with a median of 85%, resulting in a median 6.07 fold enrichment.

### DNA Extraction and Shotgun Sequencing

We extracted fecal DNA (fDNA) with the QIAGEN DNA Micro kit, following the “small volumes of blood” protocol. To improve DNA yield, we increased the lysis time to three hours, and incubated 50 µL of 56°C elution buffer on the spin column membrane for 10 minutes. DNA concentration was measured with a Qubit fluorometer. Additionally, to calculate endogenous DNA enrichment, we extracted DNA directly from five fecal samples prior to their having undergone FACS. We extracted DNA from the nine tissue and blood samples using the QIAGEN Gentra Puregene Tissue kit and DNeasy blood and tissue kit, respectively. For the fecal samples, DNA was fragmented to 350 bp with a Covaris sonicator. We built whole genome sequencing libraries with the NEB Next Ultra 2 kit using 10-11 PCR cycles. Fecal genomic libraries were sequenced on an Illumina NextSeq (2×150 PE) at the University of Calgary genome sequencing core and an Illumina HighSeq 4000 at the McDonnell Genome Institute at Washington University in St. Louis (MGI). Using ½ of one HiSeq 4000 lane, we achieved an average coverage of 12.2X across the *Cebus imitator* 1.0 genome (sample SSR-ML). Other fecal samples were sequenced to average depths of 0.1-4.4X (SI Appendix: Table S12). High-coverage (10.3-47.6X), whole genome shotgun libraries were prepared for the blood and tissue DNA samples and sequenced on an Illumina X Ten system at MGI. For population analyses within capuchins, we mapped genomic data from all 23 individuals sequenced (SI Appendix: Table S11) to the reference genome.

### Mapping and SNV Generation

We called SNVs for each sample independently using the *Cebus* genome and the GATK UnifiedGenotyper pipeline (see SI Appendix for further methodological detail). We included reads from all nine tissue/blood samples and one frozen fecal sample with high-coverage (SSR-ML). In total, we identified 4,184,363 SNVs for downstream analyses. To remove potential human contamination from sequenced libraries, we mapped trimmed reads to the *Cebus imitator* 1.0 and human (hg38) genomes simultaneously with BBsplit. Using default BBsplit parameters, we binned separately reads that mapped unambiguously to either genome. Ambiguously mapping reads (i.e. those mapping equally well to both genomes) were assigned to both genomic bins, and unmapped reads were assigned to a third bin. We calculated the amount of human genomic contamination as the percentage of total reads unambiguously mapping to the human genome (SI Appendix: Table S11). After removing contaminant reads, all libraries with at least 0.5X genomic coverage were used for population structure analysis.

In order to test the effect of fecalFACS on mapping rates, we selected five samples at random (SSR-CH, SSR-NM, SSR-LE, SSR-PR, SSR-SN) to compare pre- and post-FACS mapping rates. To test for an increase in mapping percentage, we ran a one-sample paired Wilcoxon signed-rank test on the percentages of reads that mapped exclusively to the *Cebus* genome before and after FACS. Additionally, we ran Pearson’s product moment correlations to test for an effect of the number of cells (log10 transformed) on rates of mapping, read duplication, and nanograms of input DNA. The above tests were all performed in R.

### High-coverage fecal genome comparison

We made several comparisons between our high-coverage feces-derived genome and the blood/tissue-derived genomes using window-based approaches. For each test, the feces-derived genome should fall within the range of variation for members of its population of origin (SSR). Deviations from this, for example all fecal genomes clustering together, would indicate biases in our DNA isolation methods. To assess this, we constructed 10 KB windows with a 4KB slide along the largest scaffold (21,314,911 bp) in the *C. imitator* reference genome. From these windows, we constructed plots of coverage density and the distribution of window coverage along the scaffold. Secondly, we assessed the level of heterozygosity in 1 MB / 200 KB sliding windows throughout the ten largest scaffolds. For each high-coverage genome, we plotted the density distribution of window heterozygosity. We measured genome-wide GC content with the Picard Tools CollectGcBiasMetrics function. The percentage of GC content was assessed against the distribution of normalized coverage and the number of reads in 100 bp windows per the number reads aligned to the windows.

### Population Structure

Given the large degree of difference in coverage among our samples, (less than 1X to greater than 50X), we performed pseudodiploid allele calling on all samples. For each library, at each position in the SNV set, we selected a single, random read from the sequenced library. From that read, we called the variant information at the respective SNV site for the given library. In doing so, we generated a VCF with a representative degree of variation and error for all samples.

To assess population structure and infer splits between northern and southern groups of Costa Rican white-faced capuchins, we constructed principal components plots with EIGENSTRAT (101) and built population trees with TreeMix (102). Because we ascertained variants predominantly with libraries that were of tissue/blood origin, we built principal components solely with SNVs from these libraries and projected the remaining fecal libraries onto the principal components. For our maximum likelihood trees, we used two outgroups (*Saimiri sciureus*, and *Cebus albifrons*), with *S. sciureus* serving as the root of the tree. Given the geographic distance and anthropogenic deforestation between northern and southern populations, we assumed no migration. To account for linkage disequilibrium, we grouped SNVs into windows of 1,000 SNVs. Population size history was inferred from the highest coverage non-reference individual, SSR-RM08 with PSMC, using default parameters (https://github.com/lh3/psmc).

### Local adaptation, F_ST_, heterozygosity, relatedness

For all analyses of local adaptation and heterozygosity between populations, we excluded individuals from our low-coverage dataset. We tested for the degree of relatedness among all high and low-coverage individuals using READ (103) and identified two of the high-coverage individuals from SSR, SSR-ML and SSR-CR, as potential first degree relatives (SI Appendix: Figure S10, Table S14). For all statistical analyses of high-coverage samples, we removed SSR-ML, because SSR-CR was sequenced to higher average depth.

For each individual, we calculated heterozygosity in 1 Mb / 200 Kb sliding windows across the genome for all scaffolds at least 1 Mb in length. Windows were generated with BedTools windowMaker (104) and heterozygosity was calculated as the per-site average within each window. Based upon a visual inspection of the average heterozygosity values across the genome (Figure 6), we classified a window as part of a run of homozygosity if the window’s average heterozygosity fell below 0.0002. Descriptive statistics and two-sided Wilcoxon tests were calculated in R.

For each high-coverage sample, we calculated the Hudson’s F_ST_ ratio of averages (105) in 20 kb windows with a slide of 4 Kb across the genome. Among the genes present in each window in the top 0.5% and top 0.1% of F_ST_ values, we searched for SNPs with high or moderate effects using SnpEff and identified those SNPs with high F_ST_ values (> 0.75) using VCFtools. We searched for functional enrichment of our population gene set using ToppFun in the ToppGene Suite (45). In an effort to identify candidate genes for further investigation, we set low thresholds for significance (FDR p-value < 0.1 and the minimum number of genes per category to 1).

### Chemosensory genes

The chemosensory behaviors of capuchins have been well-studied (73, 75), and taste and olfaction are suspected to play important roles in their foraging ecology. 614 orthologous olfactory receptor genes were identified in the *Cebus* reference genome using previously described methods (106). Briefly, putative OR sequences were identified by conducting TBLASTN searches against the capuchin reference assembly using functional human OR protein sequences as queries with an e-value threshold of 1e-20. For each reference-derived OR gene, we added 500 bp of flanking sequence to both the 5’ and 3’ ends in a bed file. For each individual, we extracted the OR gene region from the gVCF and generated a consensus sequence defaulting to the reference allele at variable site using bcftools (107). The number of intact, truncated, and pseudogenized OR genes in each individual were identified using the ORA pipeline (108). We considered an OR gene to be putatively functional if its amino acid sequence was at least 300 amino acids in length. ORA was further used to classify each OR into the appropriate class and subfamily with an e-value cutoff of 1e-10 to identify the functional OR subgenome for each individual (108). Taste and vomeronasal receptor genes were identified in the NCBI genome annotation, and variable positions were located by scanning the VCF with VCFtools and bash. *Cebus* opsin tuning site positions have been identified previously (68). With the high-coverage dataset, we identified the allele(s) present at each locus in the VCF. For each low-coverage fecal-derived genome, we located the position of the tuning site in the bam file using SAMtools tview and manually called the variant when possible.

## Supporting information

Supplemental Data - Gene Enrichment tables

## DECLARATIONS

### Ethics approval and consent to participate

This research adhered to the laws of Costa Rica, the United States, and Canada and complied with protocols approved by the Área de Conservación Guanacaste and by the Canada Research Council for Animal Care through the University of Calgary’s Life and Environmental Care Committee (ACC protocol AC15-0161). Samples were collected with permission from the Área de Conservación Guanacaste (ACG-PI-033-2016) and CONAGEBIO (R-025-2014-OT-CONAGEBIO; R-002-2020-OT-CONAGEBIO). Samples were exported from Costa Rica under permits from CITES and Area de Conservacion Guanacaste (2016-CR2392/SJ #S 2477, 2016-CR2393/SJ #S 2477, DGVS-030-2016-ACG-PI-002-2016; 012706) and imported with permission from the Canadian Food and Inspection Agency (A-2016-03992-4).

### Availability of data and materials

The reference genome is available at NCBI through BioProjects PRJNA298580 and PRJNA328123. RNAseq reads used in genome annotation can be accessed through PRJNA319062. The sequencing reads used in the local adaptation will be released by NCBI upon publication (PRJNA610850), and are available immediately to reviewers and editors upon request from the corresponding author.

## Acknowledgements

We thank R. Blanco Segura and M. M. Chavarria and staff from the Área de Conservación Guanacaste and Ministerio de Ambiente y Energía, and Jose Alfredo Hernández Ugalde and Angela González Grau from CONAGEBIO in Costa Rica. From Kids Saving the Rainforest, we thank the administration, volunteers and veterinarians for their help, especially Jennfier Rice and Chip Braman. For assistance in the lab, we acknowledge Kelly Kries, Gwen Duytschaever, Eva Garrett, Jene Weatherhead, Laurie Kennedy, and Yiping Liu. We thank Aoife Doherty and Mengjia Li for exploratory analyses of selection in aging-related genes and acknowledge Patrick Minx and Kim Kyung for assistance with bioinformatics. We thank Eva Wikberg, Fernando Campos, Kathy Jack, Linda Fedigan and many students, research assistants and volunteers for their contributions to the PACE database and Santa Rosa project. We also thank Jessica Lynch and Hazel Byrne for sharing comparative data. We are grateful to R. Gregory in the Centre for Genomic Research and I.C. Smith in the Advanced Research Computing at the University of Liverpool for the access to the computing resources and to Dr. J.C Opazo for insightful discussions.

## REFERENCES

1. A. R. DeCasien, S. A. Williams, J. P. Higham, Primate brain size is predicted by diet but not sociality. Nature Ecology & Evolution 1, 0112 (2017).

2. S. E. Street, A. F. Navarrete, S. M. Reader, K. N. Laland, Coevolution of cultural intelligence, extended life history, sociality, and brain size in primates. Proc. Natl. Acad. Sci. U. S. A. 114, 7908–7914 (2017).

3. S. L. Washburn, Tools and human evolution. Sci. Am. 203, 63–75 (1960).

4. H. Kaplan, K. Hill, J. Lancaster, A. Magdalena Hurtado, A theory of human life history evolution: Diet, intelligence, and longevity. Evolutionary Anthropology: Issues, News, and Reviews 9, 156–185 (2000).

5. A. D. Melin, H. C. Young, K. N. Mosdossy, L. M. Fedigan, Seasonality, extractive foraging and the evolution of primate sensorimotor intelligence. J. Hum. Evol. 71, 77–86 (2014).

6. D. M. Fragaszy, E. Visalberghi, L. M. Fedigan, The Complete Capuchin: The Biology of the Genus Cebus (Cambridge University Press, 2004).

7. Melin AD, Hogan JD, Campos FA, Wikberg E, King-Bailey G, Webb SE, Kalbitzer U, Asensio N, Murillo-Chacon E, Cheves Hernandez S, Guadamuz Chavarria A, Schaffner C, Kawamura S, Aureli F, Fedigan LM, Jack KM, Primate life history, social dynamics, ecology, and conservation: contributions from long-term research in the Área de Conservación Guanacaste, Costa Rica. Biotropica (2020) https:/doi.org/10.1111/btp.12867.

8. C. van Schaik, K. Isler, “Life-history evolution in primates” in The Evolution of Primate Societies, J. C. Mitani, J. Call, P. M. Kappeler, R. A. Palombit, J. B. Silk, Eds. (University of Chicago Press Chicago, 2012).

9. F. De Petrillo, A. G. Rosati, Ecological rationality: Convergent decision-making in apes and capuchins. Behav. Processes 164, 201–213 (2019).

10. B. J. Barrett, et al., Habitual stone-tool-aided extractive foraging in white-faced capuchins, Cebus capucinus. R. Soc. open sci. 5, 181002 (2018).

11. N. J. Emery, “Are Corvids ‘Feathered Apes’? Cognitive evolution in crows, jays, rooks and jackdaws” in Comparative Analysis of Minds, S. Watanabe, Ed. (Keio University Press: Tokyo., 2004), pp. 181–213.

12. L. Marino, Convergence of complex cognitive abilities in cetaceans and primates. Brain Behav. Evol. 59, 21–32 (2002).

13. G. Muntané, et al., Biological Processes Modulating Longevity across Primates: A Phylogenetic Genome-Phenome Analysis. Mol. Biol. Evol. 35, 1990–2004 (2018).

14. Y. Li, J. P. de Magalhães, Accelerated protein evolution analysis reveals genes and pathways associated with the evolution of mammalian longevity. Age 35, 301–314 (2013).

15. F. A. Campos, et al., Differential impact of severe drought on infant mortality in two sympatric neotropical primates. Royal Society Open Science 7, 200302 (2020).

16. M. L. Bergstrom, M. Emery Thompson, A. D. Melin, L. M. Fedigan, Using urinary parameters to estimate seasonal variation in the physical condition of female white-faced capuchin monkeys (Cebus capucinus imitator). Am. J. Phys. Anthropol. 163, 707–715 (2017).

17. M. Kuehn, H. Welsch, T. Zahnert, T. Hummel, Changes of pressure and humidity affect olfactory function. Eur. Arch. Otorhinolaryngol. 265, 299–302 (2008).

18. J. D. Mollon, “Tho’she kneel’d in that place where they grew…” The uses and origins of primate colour vision. J. Exp. Biol. (1989).

19. F. A. Campos, L. M. Fedigan, Behavioral adaptations to heat stress and water scarcity in white-faced capuchins (Cebus capucinus) in Santa Rosa National Park, Costa Rica. Am. J. Phys. Anthropol. 138, 101–111 (2009).

20. T. Proffitt, et al., Wild monkeys flake stone tools. Nature 539, 85–88 (2016).

21. P. Izar, et al., Flexible and conservative features of social systems in tufted capuchin monkeys: comparing the socioecology of Sapajus libidinosus and Sapajus nigritus. Am. J. Primatol. 74, 315–331 (2012).

22. J. R. Merkt, C. R. Taylor, “Metabolic switch” for desert survival. Proceedings of the National Academy of Sciences 91, 12313–12316 (1994).

23. K. Schmidt-Nielsen, B. Schmidt-Nielsen, Water metabolism of desert mammals 1. Physiol. Rev. 32, 135–166 (1952).

24. J. W. Cain III, P. R. Krausman, S. S. Rosenstock, J. C. Turner, Mechanisms of Thermoregulation and Water Balance in Desert Ungulates. Wildl. Soc. Bull. 34, 570–581 (2006).

25. A. Estrada, et al., Impending extinction crisis of the world’s primates: Why primates matter. Sci Adv 3, e1600946 (2017).

26. G. H. Perry, J. C. Marioni, P. Melsted, Y. Gilad, Genomic-scale capture and sequencing of endogenous DNA from feces. Mol. Ecol. 19, 5332–5344 (2010).

27. N. Snyder-Mackler, et al., Efficient Genome-Wide Sequencing and Low-Coverage Pedigree Analysis from Noninvasively Collected Samples. Genetics 203, 699–714 (2016).

28. K. L. Chiou, C. M. Bergey, Methylation-based enrichment facilitates low-cost, noninvasive genomic scale sequencing of populations from feces. Sci. Rep. 8, 1975 (2018).

29. J. Hernandez-Rodriguez, et al., The impact of endogenous content, replicates and pooling on genome capture from faecal samples. Mol. Ecol. Resour. 18, 319–333 (2018).

30. H. Long, K. Huang, Transport of Ciliary Membrane Proteins. Front Cell Dev Biol 7, 381 (2020).

31. Z. Lu, F. Wang, M. Liang, SerpinC1/Antithrombin III in kidney-related diseases. Clin. Sci. 131, 823–831 (2017).

32. S. O. Lau, J. Y. Tkachuck, D. K. Hasegawa, J. R. Edson, Plasminogen and antithrombin III deficiencies in the childhood nephrotic syndrome associated with plasminogenuria and antithrombinuria. J. Pediatr. 96, 390–392 (1980).

33. A. Citak, S. Emre, A. Sirin, I. Bilge, A. Nayır, Hemostatic problems and thromboembolic complications in nephrotic children. Pediatr. Nephrol. 14, 138–142 (2000).

34. M. Liang, et al., Molecular networks in Dahl salt-sensitive hypertension based on transcriptome analysis of a panel of consomic rats. Physiol. Genomics 34, 54–64 (2008).

35. A. P. Morris, et al., Trans-ethnic kidney function association study reveals putative causal genes and effects on kidney-specific disease aetiologies. Nat. Commun. 10, 29 (2019).

36. Y. Okada, et al., Meta-analysis identifies multiple loci associated with kidney function- related traits in east Asian populations. Nat. Genet. 44, 904–909 (2012).

37. S. Cirak, et al., ISPD gene mutations are a common cause of congenital and limb-girdle muscular dystrophies. Brain 136, 269–281 (2013).

38. T. Roscioli, et al., Mutations in ISPD cause Walker-Warburg syndrome and defective glycosylation of α-dystroglycan. Nat. Genet. 44, 581–585 (2012).

39. U. Mayer, et al., Absence of integrin alpha 7 causes a novel form of muscular dystrophy. Nat. Genet. 17, 318–323 (1997).

40. Q. Zhang, et al., Nesprin-1 and -2 are involved in the pathogenesis of Emery Dreifuss muscular dystrophy and are critical for nuclear envelope integrity. Hum. Mol. Genet. 16, 2816–2833 (2007).

41. J. A. Doe, et al., Transgenic overexpression of the α7 integrin reduces muscle pathology and improves viability in the dy(W) mouse model of merosin-deficient congenital muscular dystrophy type 1A. J. Cell Sci. 124, 2287–2297 (2011).

42. S. Amin, et al., Discovery of a drug candidate for GLIS3-associated diabetes. Nat. Commun. 9, 2681 (2018).

43. X.-A. Wang, et al., Interferon regulatory factor 9 protects against hepatic insulin resistance and steatosis in male mice. Hepatology 58, 603–616 (2013).

44. W. M. Gerding, et al., Ccdc66 null mutation causes retinal degeneration and dysfunction. Hum. Mol. Genet. 20, 3620–3631 (2011).

45. J. Chen, E. E. Bardes, B. J. Aronow, A. G. Jegga, ToppGene Suite for gene list enrichment analysis and candidate gene prioritization. Nucleic Acids Res. 37, W305–11 (2009).

46. M. M. Memon, et al., A novel WDR62 mutation causes primary microcephaly in a Pakistani family. Mol. Biol. Rep. 40, 591–595 (2013).

47. P. Stankiewicz, et al., Haploinsufficiency of the Chromatin Remodeler BPTF Causes Syndromic Developmental and Speech Delay, Postnatal Microcephaly, and Dysmorphic Features. Am. J. Hum. Genet. 101, 503–515 (2017).

48. J. Guo, et al., Developmental disruptions underlying brain abnormalities in ciliopathies. Nat. Commun. 6, 7857 (2015).

49. A. Fujita, et al., Homozygous splicing mutation in NUP13 3 causes Galloway-Mowat syndrome. Annals of Neurology 84, 814–828 (2018).

50. P. B. Crino, mTOR signaling in epilepsy: insights from malformations of cortical development. Cold Spring Harb. Perspect. Med. 5 (2015).

51. C. Bertipaglia, J. C. Gonçalves, R. B. Vallee, Nuclear migration in mammalian brain development. Semin. Cell Dev. Biol. 82, 57–66 (2018).

52. F. Laumonnier, et al., Mutations in PHF8 are associated with X linked mental retardation and cleft lip/cleft palate. J. Med. Genet. 42, 780–786 (2005).

53. R. Tacutu, et al., Human Ageing Genomic Resources: new and updated databases. Nucleic Acids Res. 46, D1083–D1090 (2018).

54. N. Fujii, et al., Sterol regulatory element-binding protein-1c orchestrates metabolic remodeling of white adipose tissue by caloric restriction. Aging Cell 16, 508–517 (2017).

55. M. Plank, D. Wuttke, S. van Dam, S. A. Clarke, J. P. de Magalhães, A meta-analysis of caloric restriction gene expression profiles to infer common signatures and regulatory mechanisms. Mol. Biosyst. 8, 1339–1349 (2012).

56. C. J. Kenyon, The genetics of ageing. Nature 464, 504–512 (2010).

57. S. Zheng, et al., Deletion of the huntingtin polyglutamine stretch enhances neuronal autophagy and longevity in mice. PLoS Genet. 6, e1000838 (2010).

58. M. Cyranka, et al., Abcc5 Knockout Mice Have Lower Fat Mass and Increased Levels of Circulating GLP-1. Obesity 27, 1292–1304 (2019).

59. V. C. Myers, M. S. Fine, The creatine content of muscle under normal conditions. Its relation to the urinary creatinine. Proc. Soc. Exp. Biol. Med. 10, 10–11 (1912).

60. J. D. Orkin, et al., Seasonality of the gut microbiota of free-ranging white-faced capuchins in a tropical dry forest. ISME J. 13, 183–196 (2019).

61. J. D. Orkin, S. E. Webb, A. D. Melin, Small to modest impact of social group on the gut microbiome of wild Costa Rican capuchins in a seasonal forest. Am. J. Primatol., e22985 (2019).

62. E. K. Mallott, K. R. Amato, P. A. Garber, R. S. Malhi, Influence of fruit and invertebrate consumption on the gut microbiota of wild white-faced capuchins (Cebus capucinus). Am. J. Phys. Anthropol. (2018) https:/doi.org/10.1002/ajpa.23395.

63. R. A. Young, Fat, Energy and Mammalian Survival. Integr. Comp. Biol. 16, 699–710 (1976).

64. M. Lopez, et al., Genomic Evidence for Local Adaptation of Hunter-Gatherers to the African Rainforest. Curr. Biol. 29, 2926–2935.e4 (2019).

65. H. Shi, et al., TLR4 links innate immunity and fatty acid-induced insulin resistance. J. Clin. Invest. 116, 3015–3025 (2006).

66. A. D. Melin, et al., The Behavioral Ecology of Color Vision: Considering Fruit Conspicuity, Detection Distance and Dietary Importance. Int. J. Primatol. 35, 258–287 (2014).

67. G. H. Jacobs, A perspective on color vision in platyrrhine monkeys. Vision Res. 38, 3307– 3313 (1998).

68. C. Hiramatsu, et al., Color-vision polymorphism in wild capuchins (Cebus capucinus) and spider monkeys (Ateles geoffroyi) in Costa Rica. Am. J. Primatol. 67, 447–461 (2005).

69. G. H. Jacobs, Evolution of colour vision in mammals. Philos. Trans. R. Soc. Lond. B Biol. Sci. 364, 2957–2967 (2009).

70. S. Kawamura, A. D. Melin, “Evolution of genes for color vision and the chemical senses in primates” in Evolution of the Human Genome I: The Genome and Gene, N. Saitou, Ed. (Springer Japan, 2018), pp. 1–59.

71. T. Hiwatashi, et al., An explicit signature of balancing selection for color-vision variation in new world monkeys. Mol. Biol. Evol. 27, 453–464 (2010).

72. S. Schreiber, et al., Neurodegeneration in the olfactory bulb and olfactory deficits in the Ccdc66 -/- mouse model for retinal degeneration. IBRO Reports 5, 43–53 (2018).

73. Y. Niimura, A. Matsui, K. Touhara, Acceleration of Olfactory Receptor Gene Loss in Primate Evolution: Possible Link to Anatomical Change in Sensory Systems and Dietary Transition. Mol. Biol. Evol. 35, 1437–1450 (2018).

74. D. Li, J. Zhang, Diet shapes the evolution of the vertebrate bitter taste receptor gene repertoire. Mol. Biol. Evol. 31, 303–309 (2014).

75. A. D. Yoder, P. A. Larsen, The molecular evolutionary dynamics of the vomeronasal receptor (class 1) genes in primates: a gene family on the verge of a functional breakdown. Front. Neuroanat. 8, 153 (2014).

76. K. Moriya-Ito, T. Hayakawa, H. Suzuki, K. Hagino-Yamagishi, M. Nikaido, Evolution of vomeronasal receptor 1 (V1R) genes in the common marmoset (Callithrix jacchus). Gene 642, 343–353 (2018).

77. S. Perry, Social traditions and social learning in capuchin monkeys (Cebus). Philos. Trans. R. Soc. Lond. B Biol. Sci. 366, 988–996 (2011).

78. K. Tsutsui, et al., Variation in ligand responses of the bitter taste receptors TAS2R1 and TAS2R4 among New World monkeys. BMC Evol. Biol. 16, 208 (2016).

79. S. M. Reader, K. N. Laland, Social intelligence, innovation, and enhanced brain size in primates. Proc. Natl. Acad. Sci. U. S. A. 99, 4436–4441 (2002).

80. A. A. Freitas, J. P. de Magalhães, A review and appraisal of the DNA damage theory of ageing. Mutat. Res. 728, 12–22 (2011).

81. K. Grube, A. Bürkle, Poly(ADP-ribose) polymerase activity in mononuclear leukocytes of 13 mammalian species correlates with species-specific life span. Proc. Natl. Acad. Sci. U. S. A. 89, 11759–11763 (1992).

82. M. Cornu, V. Albert, M. N. Hall, mTOR in aging, metabolism, and cancer. Curr. Opin. Genet. Dev. 23, 53–62 (2013).

83. J. P. de Magalhães, M. Stevens, D. Thornton, The Business of Anti-Aging Science. Trends Biotechnol. 35, 1062–1073 (2017).

84. G. Zhang, et al., Comparative analysis of bat genomes provides insight into the evolution of flight and immunity. Science 339, 456–460 (2013).

85. M. Keane, et al., Insights into the evolution of longevity from the bowhead whale genome. Cell Rep. 10, 112–122 (2015).

86. D. R. Valenzano, et al., The African Turquoise Killifish Genome Provides Insights into Evolution and Genetic Architecture of Lifespan. Cell 163, 1539–1554 (2015).

87. J. P. de Magalhães, J. Costa, G. M. Church, An analysis of the relationship between metabolism, developmental schedules, and longevity using phylogenetic independent contrasts. J. Gerontol. A Biol. Sci. Med. Sci. 62, 149–160 (2007).

88. S. Kumar, G. Stecher, M. Suleski, S. B. Hedges, TimeTree: A resource for timelines, timetrees, and divergence times. Mol. Biol. Evol. 34, 1812–1819 (2017).

89. F. Schwarz, et al., Siglec receptors impact mammalian lifespan by modulating oxidative stress. Elife 4 (2015).

90. M. A. Dyer, et al., Developmental sources of conservation and variation in the evolution of the primate eye. Proc. Natl. Acad. Sci. U. S. A. 106, 8963–8968 (2009).

91. J. W. Lynch Alfaro, et al., Explosive Pleistocene range expansion leads to widespread Amazonian sympatry between robust and gracile capuchin monkeys: Biogeography of Neotropical capuchin monkeys. J. Biogeogr. 39, 272–288 (2012).

92. J. P. Boubli, A. B. Rylands, I. P. Farias, M. E. Alfaro, J. L. Alfaro, Cebus phylogenetic relationships: a preliminary reassessment of the diversity of the untufted capuchin monkeys. Am. J. Primatol. 74, 381–393 (2012).

93. L. Fedigan, Vertebrate predation in Cebus capucinus: meat eating in a Neotropical monkey. Folia Primatol. 54, 196–205 (1990).

94. B. Bushnell, BBTools software package. http://sourceforge.net/projects/bbmap (2014).

95. D. Janzen, “Tropical dry forest: area de Conservación Guanacaste, northwestern Costa Rica” in Handbook of Ecological Restoration: Restoration in Practice, D. A. Perrow M, Ed. (Cambridge University Press, Cambridge, 2002), pp. 559–583.

96. F. A. Campos, “A synthesis of long-term environmental change in Santa Rosa” in Primate Life Histories, Sex Roles, and Adaptability - Essays in Honour of Linda M. Fedigan., Developments in Primatology: Progress and Prospects., U. Kalbitzer, K. M. Jack, Eds. (Springer, New York, 2018).

97. L. Fedigan, L. Rose-Wiles, “See how they grow: Tracking capuchin monkey populations in a regenerating Costa Rican dry forest” in Adaptive Radiations of Neotropical Primates, M. Norconk, A. L. Rosenberger, P. A. Garber, Eds. (Springer, 1996), pp. 289–307.

98. C. Rinke, et al., Obtaining genomes from uncultivated environmental microorganisms using FACS-based single-cell genomics. Nat. Protoc. 9, 1038–1048 (2014).

99. A. Morgulis, E. M. Gertz, A. A. Schäffer, R. Agarwala, WindowMasker: window-based masker for sequenced genomes. Bioinformatics 22, 134–141 (2006).

100. E. S. Rice, R. E. Green, New Approaches for Genome Assembly and Scaffolding. Annu Rev Anim Biosci 7, 17–40 (2019).

101. A. L. Price, et al., Principal components analysis corrects for stratification in genome- wide association studies. Nat. Genet. 38, 904–909 (2006).

102. J. K. Pickrell, J. K. Pritchard, Inference of population splits and mixtures from genome- wide allele frequency data. PLoS Genet. 8, e1002967 (2012).

103. J. M. Monroy Kuhn, M. Jakobsson, T. Günther, Estimating genetic kin relationships in prehistoric populations. PLoS One 13, e0195491 (2018).

104. A. R. Quinlan, I. M. Hall, BEDTools: a flexible suite of utilities for comparing genomic features. Bioinformatics 26, 841–842 (2010).

105. G. Bhatia, N. Patterson, S. Sankararaman, A. L. Price, Estimating and interpreting FST: the impact of rare variants. Genome Res. 23, 1514–1521 (2013).

106. Y. Niimura, M. Nei, Extensive gains and losses of olfactory receptor genes in mammalian evolution. PLoS One 2, e708 (2007).

107. H. Li, A statistical framework for SNP calling, mutation discovery, association mapping and population genetical parameter estimation from sequencing data. Bioinformatics 27, 2987–2993 (2011).

108. S. Hayden, et al., A cluster of olfactory receptor genes linked to frugivory in bats. Mol. Biol. Evol. 31, 917–927 (2014).

