## Supplemental appendix - tables and figures for "The evolution of ecological flexibility, large brains, and long lives: capuchin monkey genomics revealed with fecalFACS"

**This PDF file includes:**

Supplementary text
Tables S1 -S10
Figures S1 -14
FecalFACS laboratory protocol

#### SUPPLEMENTAL METHODS

*Genome-wide sequencing, genome assembly and gene annotation*

We assembled a reference genome for *Cebus imitator* from DNA extracted from the kidney of a male Costa Rican individual (KSTR64) using a short read approach (Illumina HiSeq 2500). Based on a genome estimate of 3 Gb, the total sequencing depth generated was 81X, including 50X of overlapping read-pairs (200 bp insert), 26X and 5X of 3 and 8 kbs insert read pairs, respectively. The combined sequence reads were filtered and assembled using default parameter settings with ALLPATHS-LG (1). To improve the quality of gene annotation, we isolated total RNA from the whole blood of an adult male white-faced capuchin (ID: CNS-HE) permanently residing at the KSTR wildlife rehabilitation center. The blood was immediately stored in a PAXgene blood RNA tube (Qiagen), and frozen at ultralow temperatures for subsequent use. To extract total RNA, we used the PAXgene Blood RNA kit following the manufacturer's recommended protocols. A RiboZero library construction protocol was followed according to the manufacturer's specifications and sequenced on an Illumina HiSeq 2000 instrument creating 150 bp paired-end reads. We assembled the FASTQ sequence files into transcripts with Trinity, and submitted the assembled transcriptome to the National Center for Biotechnology Information (NCBI) to assist in gene annotation. The capuchin genome assembly was annotated with the NCBI pipeline previously described here: (http://www.ncbi.nlm.nih.gov/books/NBK169439/).

Our reference genome assembly for *Cebus imitator* is composed of 7,742 scaffolds (including single contig scaffolds) with an N50 scaffold length of 5.2 Mb and an N50 contig length of 41 kb. The final ungapped assembly length is 2.6 Gb (GenBank accession: GCA_001604975.1). Our estimate of total interspersed repeats using WindowMasker (2) output is 45.8%. The numbers of annotated genes are 20,740 and 9,556 for protein-coding and non-coding genes, respectively (SI Appendix: Table S1). Measures of gene representation using the known human RefSeq set of 56,230 transcripts show an average of >94% coverage with a mean identity of 92.5%. Overall, our draft assembly metrics and gene representation are consistent with other non-human primate (NHP) short-read reference assemblies (3).

*Mapping and SNV Generation*

Reads were trimmed of sequencing adaptors with Trimmomatic (4). Subsequently, we mapped the *Cebus* reads to the *Cebus imitator* 1.0 reference genome (GCF_001604975.1) with BWA mem (5) and removed duplicates with Picard Tools (http://broadinstitute.github.io/picard/) and SAMtools (6). We called SNVs for each sample independently using the *Cebus* genome and the GATK UnifiedGenotyper pipeline (*-out_mode EMIT_ALL_SITES*) (7). Genomic VCFs were then combined using GATK's CombineVariants restricting to positions with a depth of coverage between 3 and 100, mapping quality above 30, no reads with mapping quality zero, and variant PHRED scores above 30. Sequencing reads from one of the high-coverage fecal samples (SSR-

FL) bore a strong signature of human contamination (16%), and were thus excluded from SNV generation. We included reads from nine tissue/blood samples and one frozen fecal sample with high-coverage (SSR-ML). In total, we identified 4,184,363 SNVs for downstream analyses.

To remove potential human contamination from sequenced libraries, we mapped trimmed reads to the *Cebus imitator* 1.0 and human (hg38) genomes simultaneously with BBsplit . Using default BBsplit parameters, we binned separately reads that mapped unambiguously to either genome. Ambiguously mapping reads (i.e. those mapping equally well to both genomes) were assigned to both genomic bins, and unmapped reads were assigned to a third bin. We calculated the amount of human genomic contamination as the percentage of total reads unambiguously mapping to the human genome (SI Appendix: Table S11). After removing contaminant reads, all libraries with at least 0.5X genomic coverage were used for population structure analysis.

In order to test the effect of fecalFACS on mapping rates, we selected five samples at random (SSR-CH, SSR-NM, SSR-LE, SSR-PR, SSR-SN) to compare pre- and post-FACS mapping rates. To test for an increase in mapping percentage, we ran a one-sample paired Wilcoxon signed-rank test on the percentages of reads that mapped exclusively to the *Cebus* genome before and after FACS. Additionally, we ran Pearson's product moment correlations to test for an effect of the number of cells (log10 transformed) on rates of mapping, read duplication, and nanograms of input DNA. The above tests were all performed in R.

*Phylogenetic arrangement and data treatment*

The phylogenetic arrangement in this study included 14 species as outgroups to *C. imitator*: three Platyrrhini (*Callithrix jacchus*, *Aotus nancymaae*, *Saimiri* boliviensis), six Catarrhini (*Macaca mulatta*, *Rhinopithecus roxellana*, *Nomascus leucogenys*, *Pan troglodytes*, *Homo sapiens*, *Pongo abelii*), one Strepsirrhini (*Microcebus murinus*), one rodent (*Mus musculus*), and three Laurasiatheria (*Canis lupus familiaris*, *Bos taurus,* and *Sus scrofa*). Genomic cds were downloaded from Ensembl and NCBI (SI Appendix: Table S13). The sequences per genome were clustered using CD-HITest version 4.6 (8) with a sequence identity threshold of 90% and an alignment coverage control of 80%. To remove low quality sequences and keep the longest transcript per gene, we used TransDecoder.LongOrfs and TransDecoder.Predict (https://transdecoder.github.io) with default criteria.

*Orthology identification*

The orthology assessment was performed with OMA stand-alone v. 2.3.1 (9). The OMA algorithm makes strict pairwise "all-against-all" sequence comparisons and identifies the orthologous pairs (genes related by speciation events) based on evolutionary distances. These orthologous genes were clustered into Orthologous Groups (OGs). All OGs included one ortholog sequence from capuchin and at least one outgroup. The tree topology was obtained from TimeTree (http://www.timetree.org/ (10)). We identified 7,519 OGs present among the 15 species. Each orthogroup shared by all species was translated into amino acids using the function pxtlate -s in phyx (11). Amino acid sequences were aligned using the L-INS-i algorithm from MAFFT v.7 (12). We generated codon alignments using pxaa2cdn in phyx. To avoid false positives in low quality regions, the codon alignments were cleaned with the codon.clean.msa algorithm in rphast (13), using human as a reference sequence. We used conservative methodologies of homology and data cleaning to obtain a smaller number of orthologous genes that avoided false positives with high confidence.

We recovered 23,402 Orthologous Groups (OGs). Capuchins share 18,475 OGs with human, 17,589 OGs with rhesus macaque, 15,582 OGs with mouse, and 14,404 OGs with dog. When we included orthologous genes that are present simultaneously in all 15 species, we recovered 7,519 OGs, which we subsequently used in the natural selection analyses ($d_N/d_S=\omega$). While including a broad diversity of mammals decreases the number of OGs available for analysis, PAML benefits from including variation in its phylogenetic design. Including a greater number of species decreases the number of orthologous genes available for analyses and introduces the risk of missing some genes under positive selection. However, including a higher number of species, some of which are not closely related to *Cebus*, improves the biological and statistical background for the natural selection analysis (14). Additionally, including these species of Laurasiatheria (dog, cow, and pig) not only increased variation within the phylogenetic design, but because they are exceptionally high-quality genome assemblies, including them improves our confidence in recovering high-quality orthologous gene sequences. We identified 612 genes under positive selection ($p<0.05$ after FDR correction) in the *Cebus* lineage using the branch model. We also performed a branch-site test using codeml in PAML (15) and identified a second set of 748 genes under positive selection in *Cebus* (Supplemental Data: IDs).

*Positive natural selection analysis through codon-based models of evolution and enrichment tests*

To evaluate signals consistent with positive selection in the *C. imitator* genome, we explored variation in the ratio of non-synonymous and synonymous substitutions ($d_N/d_S=\omega$) in the ancestor of *Cebus*. We used branch and branch-site substitution models with a maximum likelihood approach in PAML v4.9 (15), which we implemented through the python framework ETE-toolkit with the ete-evol function (16). We compared the null model where the omega ($\omega$) value in the branch marked as foreground was set with 1, with the model where the $\omega$ value was estimated from the data (17). Likelihood ratio tests (LRT) were used to test for significance between the models and probability values were adjusted with a false discovery rate correction for multiple testing with a q-value $< 0.05$ for the two positive selection models (branch and branch-site).

We performed functional annotation analysis using ToppFunn in the ToppGene Suite (18) with default parameters. To ascertain which ontology processes the genes with signals of positive selection were involved, we focused the enrichment analysis on two functional categories: GO Biological Processes (BP) and the DisGeNET BeFree disease database. Finally, the genes with positive selection signal were intersected with the GenAge and CellAge databases (build 19,307 genes) (19).

#### SUPPLEMENTAL TABLES

**Table S1**: Number of annotated and coding genes from *Cebus imitator* reference genome.
https://www.ncbi.nlm.nih.gov/genome/annotation_euk/Cebus_capucinus_imitator/100/

**Table S2**: Genes falling in high $F_{ST}$ (top 0.5%) windows and those with high $F_{ST}$ (>0.75) non-synonymous SNPs.

| **SNPEFF High/Moderate Fixed SNPs: 26 genes** |
|---|
| ABCC5, CCDC66, DIAPH2, EPS15L1, FAM118A, GPRASP1, IRF9, IRS4, ISPD, ITGA7, LYPLA2, METTL10, MUC19, PCDHB1, PCDHB3, PDLIM2, RPA4, RRN3, SPOPL, STPG3, TLR4, TRIM15, WFDC10A, WLS, WRB, ZSCAN20 |
| **Top 0.5% of Fst Windows with SNPEFF High/Moderate FST (>= 0.75): 39 genes** |
| ADGRD1, ASCC3, AXDND1, BBOF1, CCDC66, DTHD1, EFCAB7, EPS15L1, ERVFRD-1, FAM118A, FAM208A, FAM227B, GLIS3, INSL6, IRF9, ISPD, ITGA7, MAMDC4, MASTL, METTL10, NCKAP5L, PCDHB1, PCDHB3, PTX4, REC114, RRN3, SERPINC1, SMIM13, SPHKAP, STPG3, SYNE2, TBC1D5, TCOF1, TMEM132C, TMEM204, VRK3, WLS, ZNF106, ZSCAN20 |
| **Top 0.5% of Fst Windows: 299 genes** |
| ABCC1, ABCD2, ABI1, ADAM32, ADAMTSL1, ADGRD1, AKT1, ALDH6A1, ANAPC13, ANKRD26, ANO4, ANTXR1, ARAP1, ARHGEF28, ASAP1, ASCC3, ATG5, ATP2B2, ATP2B3, ATP5J2, ATP8B4, AVPR1B, AWAT2, AXDND1, BBOF1, BCAS3, BEND4, BLOC1S1, BMP15, BTBD11, CABP1, CACNA1B, CACYBP, CALN1, CAPN10, CAPN3, CAPRIN1, CCBE1, CCDC170, CCDC53, CCDC66, CD74, CEP63, CHCHD3, CHMP4A, CHP2, CIB4, CKAP2, CLDN19, CLDN2, CLVS1, CNEP1R1, CNNM1, CNTNAP2, COG3, COL14A1, COQ6, CPA6, CPM, CPSF4, CSMD2, CUL4B, CUX2, CYTIP, DBX2, DCAF11, DCAF4L1, DCDC2C, DLG3, DNAH8, DPP10, DPY30, DSTYK, DTWD1, EDA, EDF1, EEF1E1, EFCAB7, EMC9, ENTPD5, ENTPD7, EPHA2, EPHB1, EPS15L1, ERVFRD-1, ESR2, ESX1, EXOC4, EXT2, FAM101A, FAM118A, FAM161B, FAM166A, FAM208A, FAM227B, FAM53B, FAM58A, FCHSD2, FER1L6, FGL1, FHL1, FITM1, FLI1, GALNT13, GALNT2, GANC, GBE1, GDPD4, GHR, GLIS3, GMPR2, GNG12, HS3ST4, IFT140, IFT74, INPP4B, INSL6, IPO4, IRF9, ISPD, ITGA7, ITGA8, ITGB3BP, JAK2, KAZN, KCNH3, KCNIP4, KCNJ5, KCNV1, KCTD3, KHDRBS2, KLHL1, KY, LAMA2, LARGE1, LIN52, LPAR4, LRP8, LRRC31, LRRC34, LRRIQ4, LYPD6B, MAMDC4, MASTL, MCRS1, MDP1, MECOM, MEMO1, METTL10, METTL7B, MPP7, MPRIP, MSI2, MTHFD1L, MTHFS, MYH14, MYO3B, NAV2, NCKAP5L, NEDD8, NEK3, NELFB, NELL1, NET1, NID1, NKX6-1, NMNAT3, NPTN, NRG1, NTM, NUP210, NUP37, NUP62CL, OGT, OTOF, OTUD7B, P2RY10, P3H1, PAQR8, PARD3B, PARPBP, PAX9, PBX1, PCDHB1, PCDHB3, PDSS1, PGBD5, PGM5, PGPEP1L, PHACTR1, PHC2, PHLDB2, PHPT1, PITPNC1, PKIB, PKP4, PLCXD2, PLGRKT, PLPPR1, PLXDC1, PLXDC2, PNO1, PPARGC1A, PPP3R1, PRMT9, PRR32, PRRC2B, PSME1, PSME2, PTX4, RABGAP1L, RABL6, RAPGEF2, RBM41, REC114, REC8, REL, RFX3, RNASE1, RNASE2, RNF128, RNF31, RNGTT, RNH1, RPL26L1, RPP21, RRAS2, RRN3, SASH1, SCAF11, SEMA3E, SERPINC1, SHANK2, SLC25A21, SLC2A13, SLC30A6, SLC30A9, SLC35D3, SLC38A4, SLC39A8, SLC44A3, SMC1B, SMIM13, SNAP23, SNTG2, SNX12, SPAST, SPHKAP, SPON1, STARD10, STPG3, SULT4A1, SYNE2, TBC1D5, TBX18, TCOF1, TELO2, TENM4, TEX11, TGM1, TINF2, TM9SF1, TMCC2, TMEM117, TMEM132C, TMEM176B, TMEM184C, TMEM252, TMEM45B, TMEM87A, TNR, TRIM36, TSSK4, TTC6, TTLL7, TUBB4B, TYROBP, UBQLN1, UBR2, UNC45B, USH2A, VRK3, WDR92, WLS, WNT4, WWC1, WWTR1, YME1L1, ZBTB37, ZBTB42, ZKSCAN5, ZNF106, ZNF385B, ZNF394, ZNF473, ZNF536, ZNF695, ZNF789, ZSCAN20 |

**Table S3:** Genes associated with diabetic nephropathy found in GWAS Catalogue and top 0.5% of Fst windows

| Diabetic Nephropathy Genes (GWAS Catalogue) |
|---|
| ABRACL, ACTR3, AFF3, AGMO, APOL1, ARL6IP1P3, AUH, B4GALT1, BCAS1, BSND, BTBD11, C19orf81, CASC9, CCDC80, CGNL1, CHAT, CNTNAP2, COL4A1, CRCP, CYGB, CYP24A1, DIAPH3, EFCAB8, ELMO1, ERBB4, FOXP4, FTO, FUT9, GABRR1, GABRR2, GPR158, GSTM5P1, GUCY1A1, HAND2, HMGB1P13, HMGB1P50, HMGN1P17, IGSF22, INSYN1, ITGA6, ITPR1, JAK1, KCNH7, KDM4D, KERA, KL, KRTAP3-2, KRTAP3-3, LAMC1, LHX3, LIMK2, LINC01003, LINC01151, LINC01191, LINC01249, LINC01276, LINC01738, LINC02237, LINC02511, LINC02607, LRP8, LSAMP, LY86, MIR646HG, MTNR1B, MYH9, NAV3, NPM1P48, NRG3, PCSK6, PRCD, PROM1, PTPN13, RAD51B, RAPGEF5, RBL1, REPS1, RN7SKP82, RN7SL72P, RN7SL865P, RNA5SP86, RNF10, RNU4-37P, RNU6-1059P, RNU6-34P, RNU6-41P, RNU7-88P, RPA2P2, RPL26P31, RPL37A, RPS12, RPSAP52, RYR3, SAMHD1, SASH1, SCGB2B2, SLC30A8, SLC7A15P, SMIM13, SORCS3, SPINK4, STEAP1B, STX8, SUCLG2, SYN2, TAPT1, TBC1D27P, TBC1D31, TBC1D5, TBXAS1, TENM2, TNFRSF19, TOMM22P3, TRABD2B, TTC21B, TTC39C, WWC1 |
| **Genes in top 0.5% of Fst Windows** |
| BTBD11, CTNAP2, LRP8, SASH1, SMIM13, TBC1D5, WWC1 |

**Table S4**: Codons present at long/medium wavelength opsin tuning sites on scaffold NW_016107914.1. Heterozygous positions cannot be known for most low-coverage samples. Empty cells indicate that zero reads covered a given low-coverage codon position.

| Individual | Population | 180 | 277 | 285 | Coverage |
|---|---|---|---|---|---|
| | | 50434-50436 | 46877-46879 | 46853-46855 | |
| CNS-HE | North | S | Y | T | High |
| SSR-CR | North | A | F | T | High |
| SSR-FG | North | | Y | | Low |
| SSR-FL | North | S | Y | T | Low |
| SSR-KI | North | S | Y | | Low |
| SSR-LU | North | A | Y | T/A | Low |
| SSR-ML | North | A | F | T | High |
| SSR-RF | North | S | | | Low |
| SSR-RM08 | North | S | Y | T | High |
| SSR-T5-1 | North | A/S | Y/F | T/A | High |
| SSR-TH | North | A | F | T/A | High |

| | | | | | |
|---|---|---|---|---|---|
| SSR-TY | North | | Y | | Low |
| KSTR116 | South | A | F | T/A | High |
| KSTR159 | South | S | Y | T | High |
| KSTR29 | South | S | Y | T | High |
| REF/KSTR64 | South | A | F | T | High |

**Table S5**: Number and category of olfactory receptor genes identified per individual

| Individual | Population | Intact | Truncated | Pseudogene | Total |
|---|---|---|---|---|---|
| CNS-HE | North | 408 | 46 | 160 | 614 |
| SSR-CR | North | 409 | 45 | 160 | 614 |
| SSR-ML | North | 408 | 45 | 161 | 614 |
| SSR-RM08 | North | 411 | 44 | 159 | 614 |
| SSR-T5-1 | North | 409 | 46 | 159 | 614 |
| SSR-TH | North | 410 | 44 | 160 | 614 |
| KSTR116 | South | 410 | 44 | 160 | 614 |
| KSTR159 | South | 410 | 45 | 159 | 614 |
| KSTR29 | South | 407 | 46 | 161 | 614 |
| REF/KSTR64 | South | 408 | 45 | 161 | 614 |

**Table S6**: Number of intact olfactory receptor genes per gene family, as identified by ORA (20).

| Individual | Population | 1 3 7 | 2 13 | 4 | 5 8 9 | 6 | 10 | 11 | 12 | 14 | 51 | 52 | 55 | 56 | Total Intact | Total OR genes |
|---|---|---|---|---|---|---|---|---|---|---|---|---|---|---|---|---|
| CNS-HE | North | 26 | 60 | 54 | 87 | 41 | 44 | 8 | 1 | 1 | 41 | 40 | 1 | 6 | 410 | 614 |
| SSR-CR | North | 25 | 60 | 53 | 89 | 40 | 44 | 8 | 1 | 1 | 41 | 40 | 1 | 6 | 409 | 614 |
| SSR-ML | North | 25 | 60 | 53 | 88 | 40 | 44 | 8 | 1 | 1 | 41 | 40 | 1 | 6 | 408 | 614 |

| | | | | | | | | | | | | | | | |
|---|---|---|---|---|---|---|---|---|---|---|---|---|---|---|---|
| SSR-RM08 | North | 26 | 60 | 54 | 90 | 41 | 44 | 8 | 1 | 1 | 41 | 40 | 1 | 6 | 413 | 614 |
| SSR-T5-1 | North | 25 | 60 | 54 | 89 | 41 | 44 | 8 | 1 | 1 | 41 | 40 | 1 | 6 | 411 | 614 |
| SSR-TH | North | 26 | 60 | 54 | 89 | 41 | 44 | 8 | 1 | 1 | 41 | 40 | 1 | 6 | 412 | 614 |
| KSTR116 | South | 24 | 60 | 54 | 88 | 40 | 44 | 8 | 1 | 1 | 41 | 41 | 1 | 6 | 409 | 614 |
| KSTR159 | South | 26 | 60 | 54 | 87 | 40 | 45 | 8 | 1 | 1 | 41 | 40 | 1 | 6 | 410 | 614 |
| KSTR29 | South | 24 | 60 | 54 | 87 | 40 | 44 | 8 | 1 | 1 | 41 | 40 | 1 | 6 | 407 | 614 |
| REF/KSTR64 | South | 25 | 60 | 54 | 87 | 40 | 44 | 8 | 1 | 1 | 41 | 40 | 1 | 6 | 408 | 614 |

**Table S7**: Taste and vomeronasal receptor genes identified in the *Cebus imitator* reference genome

| **GeneID** | **Symbol** | **Description** |
|---|---|---|
| 108282384 | TAS1R1 | taste 1 receptor member 1 |
| 108287411 | TAS1R2 | taste 1 receptor member 2 |
| 108313929 | TAS1R3 | taste 1 receptor member 3 |
| 108281399 | TAS2R1 | taste 2 receptor member 1 |
| 108295816 | TAS2R10 | taste 2 receptor member 10 |
| 108295823 | TAS2R13 | taste 2 receptor member 13 |
| 108295819 | TAS2R14 | taste 2 receptor member 14 |
| 108288651 | TAS2R16 | taste 2 receptor member 16 |
| 108283429 | TAS2R3 | taste 2 receptor member 3 |
| 108283437 | TAS2R38 | taste 2 receptor member 38 |
| 108291118 | TAS2R39 | taste 2 receptor member 39 |
| 108283430 | TAS2R4 | taste 2 receptor member 4 |
| 108291117 | TAS2R40 | taste 2 receptor member 40 |
| 108295820 | TAS2R42 | taste 2 receptor member 42 |
| 108283431 | TAS2R5 | taste 2 receptor member 5 |

| | | |
|---|---|---|
| 108291079 | TAS2R60 | taste 2 receptor member 60 |
| 108295815 | TAS2R7 | taste 2 receptor member 7 |
| 108295849 | TAS2R8 | taste 2 receptor member 8 |
| 108295818 | TAS2R9 | taste 2 receptor member 9 |
| 108282509 | LOC108282509 | taste receptor cell protein 1 |
| 108295824 | LOC108295824 | taste receptor type 2 member 13-like |
| 108295822 | LOC108295822 | taste receptor type 2 member 20-like |
| 108291100 | LOC108291100 | taste receptor type 2 member 41-like |
| 108295826 | LOC108295826 | taste receptor type 2 member 45-like |
| 108295821 | LOC108295821 | taste receptor type 2 member 46-like |
| 108295827 | LOC108295827 | taste receptor type 2 member 50 |
| 108295828 | LOC108295828 | taste receptor type 2 member 50-like |
| 108291080 | LOC108291080 | taste receptor type 2 member 62-like |
| 108289932 | LOC108289932 | vomeronasal type-1 receptor 1 |
| 108297370 | LOC108297370 | vomeronasal type-1 receptor 3-like |
| 108283963 | LOC108283963 | vomeronasal type-1 receptor 4-like |
| 108289502 | LOC108289502 | vomeronasal type-1 receptor 4-like |
| 108297824 | LOC108297824 | vomeronasal type-1 receptor 42-like |
| 108300766 | LOC108300766 | vomeronasal type-1 receptor 90 |
| 108300137 | LOC108300137 | vomeronasal type-2 receptor 1-like |

**Table S8**: Reference allele frequencies of SNPs in taste receptor genes that are fixed in the northern population but variable in the south.

| Gene | Scaffold | Site | Ref | Alt | Reference Allele Frequency | |
|---|---|---|---|---|---|---|
| | | | | | North | South |
| TAS1R | NW_016107330.1 | 9124966 | A | G | 0 | 0.625 |

| | | | | | | |
|---|---|---|---|---|---|---|
| TAS2R4 | NW_016107575.1 | 2971677 | A | G | 1 | 0.375 |

**Table S9:** Genes putatively linked to longevity as identified in the GenAge database.

| GenAge Database | | CellAge Genes | |
|---|---|---|---|
| Gene ID | Ensembl gene ID | | Gene ID |
| PARP1 | ENSG00000143799 | CSNK2A1 | ENSG00000101266 |
| MAP3K5 | ENSG00000197442 | RUVBL2 | ENSG00000183207 |
| RBBP6 | ENSG00000122257 | NEK4 | ENSG00000114904 |
| SIN3A | ENSG00000169375 | KDM5B | ENSG00000117139 |
| MYOF | ENSG00000138119 | SIN3B | ENSG00000127511 |
| POLA1 | ENSG00000101868 | SREBF1 | ENSG00000072310 |
| ZMPSTE24 | ENSG00000084073 | CDK4 | ENSG00000135446 |
| RB1 | ENSG00000139687 | PIK3C2A | ENSG00000011405 |
| INSR | ENSG00000171105 | FXR1 | ENSG00000114416 |
| PTK2B | ENSG00000120899 | RB1 | ENSG00000139687 |
| ARHGAP1 | ENSG00000175220 | MAP2K3 | ENSG00000034152 |
| NCOR1 | ENSG00000141027 | CTNNAL1 | ENSG00000119326 |
| TRRAP | ENSG00000196367 | SPOP | ENSG00000121067 |
| PI4KB | ENSG00000143393 | SMARCA4 | ENSG00000127616 |
| MAPK9 | ENSG00000050748 | ACLY | ENSG00000131473 |
| MPDZ | ENSG00000107186 | LIMK1 | ENSG00000106683 |
| HTT | ENSG00000197386 | GRK6 | ENSG00000198055 |
| MTOR | ENSG00000198793 | SMG1 | ENSG00000157106 |
| STRN3 | ENSG00000196792 | KDM4A | ENSG00000066135 |
| PCMT1 | ENSG00000120265 | TPR | ENSG00000047410 |
| MED1 | ENSG00000125686 | TXNIP | ENSG00000265972 |
| TCF3 | ENSG00000071564 | | |
| SMARCA1 | ENSG00000102038 | | |
| TRIP12 | ENSG00000153827 | | |
| FAS | ENSG00000026103 | | |
| NCOR2 | ENSG00000196498 | | |
| MDM2 | ENSG00000135679 | | |

**Table S10**: Median heterozygosity of high-coverage samples from genome wide 1 Mb windows with a 200 Kb slide.

| Population | Individual | Median Heterozygosity |
|---|---|---|
| North | CNS-HE | 0.00046994 |
| North | SSR-CR | 0.00052988 |
| North | SSR-ML | 0.00053136 |
| North | SSR-RM08 | 0.00051522 |
| North | SSR-T5-1 | 0.00053583 |
| North | SSR-TH | 0.00057165 |
| South | KSTR116 | 0.00065454 |
| South | KSTR159 | 0.00071533 |
| South | KSTR29 | 0.00067446 |
| South | KSTR64 | 0.00069666 |

**Table S11:** Origins, preservation, and average depth of coverage information for *Cebus imitator* samples. Reported coverages for the high-coverage samples (CNS-HE, SSR-CR, SSR-ML, SSR-RM08, SSR-T5-1, SSR-TH, KSTR116, KSTR159, KSTR29, KSTR64) have not been processed with BBsplit.

| Region | Individual | Sample | Site | Sample Type | Preservation | X Coverage |
|---|---|---|---|---|---|---|
| North | CNS-HE | CNS-HE | Cañas | Blood | Frozen | 10.3 |
| North | SSR-CH | SSR-CH | Sector Santa Rosa | Feces | RNA*later* | 0.4 |
| North | SSR-FG | SSR-FG | Sector Santa Rosa | Feces | RNA*later* | 2.0 |
| North | SSR-FL | SSR-FL | Sector Santa Rosa | Feces | Frozen | 4.4 |
| North | SSR-FN | SSR-FN | Sector Santa Rosa | Feces | RNA*later* | 2.8 |
| North | SSR-KI | SSR-KI | Sector Santa Rosa | Feces | RNA*later* | 1.0 |
| North | SSR-LE | SSR-LE | Sector Santa Rosa | Feces | RNA*later* | 0.3 |
| North | SSR-LU | SSR-LU | Sector Santa Rosa | Feces | RNA*later* | 2.0 |

| | | | | | | |
|---|---|---|---|---|---|---|
| North | SSR-ML | SSR-ML | Sector Santa Rosa | Feces | Frozen | 12.2 |
| North | | SSR-ML | Sector Santa Rosa | Feces | RNA*later* | 1.9 |
| North | SSR-NM | SSR-NM | Sector Santa Rosa | Feces | RNA*later* | 0.4 |
| North | SSR-PR | SSR-PR | Sector Santa Rosa | Feces | RNA*later* | 0.1 |
| North | SSR-RF | SSR-RF | Sector Santa Rosa | Feces | RNA*later* | 0.7 |
| North | SSR-SB | SSR-SB1 | Sector Santa Rosa | Feces | RNA*later* | 1.1 |
| | | SSR-SB2 | | Feces | RNA*later* | |
| North | SSR-SN | SSR-SN | Sector Santa Rosa | Feces | RNA*later* | 0.2 |
| North | SSR-TY | SSR-TY | Sector Santa Rosa | Feces | RNA*later* | 1.5 |
| North | SSR-CR | SSR-CR | Sector Santa Rosa | Ear Punch | Frozen | 30.1 |
| North | SSR-RM08 | SSR-RM08 | Sector Santa Rosa | Lung | Frozen | 47.6 |
| North | SSR-T5-1 | SSR-T5-1 | Sector Santa Rosa | Kidney | Frozen | 17.0 |
| North | SSR-TH | SSR-TH | Sector Santa Rosa | Kidney | Frozen | 20.4 |
| South | KSTR116 | KSTR116 | Manuel Antonio | Blood | Frozen | 20.3 |
| South | KSTR159 | KSTR159 | Manuel Antonio | Blood | Frozen | 16.0 |
| South | KSTR29 | KSTR29 | Manuel Antonio | Blood | Frozen | 19.2 |
| South | KSTR64 | KSTR64 | Quepos | Blood | Frozen | 19.5 |

**Table S12:** FACS and mapping results from *C. imitator* fecal samples. All coverage values are reported after BBsplit filtration.

| | | | | | **% Reads Mapping** | | | | | |
|---|---|---|---|---|---|---|---|---|---|---|
| **Monkey** | **Library** | **Cells** | **PCR Cycles** | **Total DNA (ng)** | **BWA mem** | **BBsplit *Cebus*** | **Unique *Cebus*** | **Duplicate *Cebus*** | **BBsplit Human** | **X Coverage** |
| SSR-ML | SSR-ML Frozen | 2546 | 11 | 10.50 | 96 | 90 | 85 | 5 | 1 | 11.7 |

| | | | | | | | | | | | |
|---|---|---|---|---|---|---|---|---|---|---|---|
| | SSR-ML RNAlater | 42837 | 10 | 8.26 | | 88 | 67 | 63 | 4 | 1 | 1.9 |
| SSR-FL | SSR-FL | 4405 | 12 | 6.72 | | 80 | 42 | 40 | 3 | 16 | 4.4 |
| SSR-FN | SSR-FN | 62601 | 8 | 21.50 | | 97 | 93 | 86 | 6 | 1 | 2.8 |
| SSR-FG | SSR-FG | 580 | 10 | 9.75 | | 94 | 84 | 60 | 24 | 3 | 2.0 |
| SSR-LU | SSR-LU | 8998 | 10 | 8.00 | | 93 | 84 | 72 | 11 | 1 | 2.0 |
| SSR-TY | SSR-TY | 140 | 10 | 7.70 | | 98 | 94 | 64 | 30 | 1 | 1.5 |
| SSR-SB | SSR-SB 2 | 129 | 10 | 9.00 | | 79 | 60 | 39 | 22 | 1 | 1.1 |
| | SSR-SB 1 | 11944 | 10 | 6.25 | | 55 | 11 | 8 | 2 | 1 | |
| SSR-KI | SSR-KI | 612 | 10 | 9.00 | | 93 | 78 | 45 | 33 | 6 | 1.0 |
| SSR-RF | SSR-RF | 257 | 10 | 10.00 | | 92 | 78 | 38 | 40 | 5 | 0.7 |
| SSR-NM | SSR-NM | 3336 | 11 | 3.38 | | 98 | 95 | 92 | 3 | 1 | 0.4 |
| SSR-CH | SSR-CH | 957 | 11 | 4.06 | | 93 | 85 | 80 | 5 | 1 | 0.4 |
| SSR-LE | SSR-LE | 1612 | 11 | 2.96 | | 96 | 91 | 81 | 11 | 1 | 0.3 |
| SSR-SN | SSR-SN | 1866 | 11 | 3.96 | | 92 | 79 | 74 | 6 | 3 | 0.2 |
| SSR-PR | SSR-PR | 12316 | 11 | 3.13 | | 78 | 64 | 58 | 6 | 1 | 0.1 |
| Median | | 2206 | 10 | 7.85 | | 93 | 82 | 64 | 6 | 1 | 1.1 |

**Table S13**: Reference Genomes used in PAML analysis

| Species | Common Name | Center | Assembly Name | GenBank Assembly ID | Year | Coding Genes | Scaffolds | Scaffold N50 Mb | Contigs | Contig N50 | Hyperlink |
|---|---|---|---|---|---|---|---|---|---|---|---|
| *Aotus nancymaae* | Ma's night monkey | Baylor | Anan_2.0 | GCA_000952055.2 | 2017 | 20412 | 28,922 | 8 | 112,851 | 126Kb | https://www.ncbi.nlm.nih.gov/assembly/GCF_000952055.2/ |
| *Bos taurus* | cow | USDA | ARS-UCD1.2 | GCA_002263795.2 | 2018 | 21867 | 2,211 | 103 | 2,597 | 25Mb | https://www.ncbi.nlm.nih.gov/assembly/GCF_002263795.1/ |
| *Callithrix jacchus* | marmoset | Broad Institute | ASM275486v1 | GCA_002754865.1 | 2017 | 19690 | 39,944 | 129 | 88,439 | 155Kb | https://www.ncbi.nlm.nih.gov/assembly/GCA_002754865.1/ |
| *Canis lupus familiaris* | dog | Broad Institute | CanFam3.1 | GCA_000002285.2 | 2011 | 19856 | 3,310 | 45 | 27,106 | 267Kb | https://www.ncbi.nlm.nih.gov/assembly/GCF_000002285.3/ |
| *Cebus capucinus imitator* | capuchin | MGI | Cebus_imitator-1.0 | GCA_001604975.1 | 2016 | 20317 | 7,156 | 5 | 140,597 | 41Kb | https://www.ncbi.nlm.nih.gov/assembly/GCF_001604975.1/ |
| *Homo sapiens* | human | GRC | GRCh38.p12 | GCA_000001405.27 | 2017 | 20418 | 472 | 67 | 998 | 57Mb | https://www.ncbi.nlm.nih.gov/assembly/GCF_000001405.38/ |
| *Macaca mulatta* | rhesus macaque | Baylor | Mmul_8.0.1 | GCA_000772875.3 | 2015 | 21099 | 286,263 | 4 | 348,494 | 107Kb | https://www.ncbi.nlm.nih.gov/assembly/GCF_000772875.2/ |
| *Microcebus murinus* | mouse lemur | Baylor | Mmur_3.0 | GCA_000165445.3 | 2017 | 18895 | 7,678 | 108 | 50,984 | 210Kb | https://www.ncbi.nlm.nih.gov/assembly/GCF_000165445.2/ |
| *Mus musculus* | mouse | GRC | GRCm38.p6 | GCA_000001635.8 | 2017 | 22600 | 162 | 54 | 605 | 32Mb | https://www.ncbi.nlm.nih.gov/assembly/GCF_000001635.26/ |
| *Nomascus leucogenys* | gibbon | Baylor/Broad/MGI | Nleu_3.0 | GCA_000146795.3 | 2012 | 20794 | 17,524 | 52 | 197,900 | 35Kb | https://www.ncbi.nlm.nih.gov/assembly/GCF_000146795.2/ |
| *Pan troglodytes* | chimpanzee | UW | Clint_PTRv2 | GCA_002880755.3 | 2018 | - | 4,432 | 53 | 5,061 | 12Mb | https://www.ncbi.nlm.nih.gov/assembly/GCF_002880755.1/ |
| *Pongo abelii* | orangutan | UW | Susie_PABv2 | GCA_002880775.3 | 2018 | - | 5,300 | 98 | 5,814 | 11Mb | https://www.ncbi.nlm.nih.gov/assembly/GCF_002880775.1 |
| *Rhinopithecus roxellana* | golden snub-nosed monkey | Novogene | Rrox_v1 | GCA_000769185.1 | 2017 | 21289 | 135,512 | 1.5 | 196,797 | 77Kb | https://www.ncbi.nlm.nih.gov/assembly/GCF_000769185.1/ |
| *Saimiri boliviensis boliviensis* | Bolivian squirrel monkey | Broad Institute | SaiBol1.0 | GCA_000235385.1 | 2011 | 19380 | 2,686 | 18 | 151,414 | 38Kb | https://www.ncbi.nlm.nih.gov/assembly/GCF_000235385.1/ |
| *Sus scrofa* | pig | SGSC | Sscrofa11.1 | GCA_000003025.6 | 2017 | 22452 | 706 | 88 | 1,118 | 48Mb | https://www.ncbi.nlm.nih.gov/assembly/GCF_000003025.6/ |

**Table S14:** Pairwise estimates of relatedness generated from READ (21).

| Pair Individuals | Relationship | Z_upper | Z_lower |
|---|---|---|---|
| SSR-ML SSR-CR | First Degree | 0.32102927 | -25.612329 |
| SSH-FG SSH-LE | First Degree | 3.2233821 | -23.53498 |
| SSH-LE SSH-PR | First Degree | 4.74777227 | -16.536803 |
| SSH-PR SSH-SB | First Degree | 5.8023088 | -19.829098 |
| SSH-FL SSH-RF | First Degree | 8.78614775 | -21.784945 |
| SSH-FL SSH-FG | Second Degree | 1.33327833 | -12.877227 |
| SSH-FG SSH-PR | Second Degree | 2.01585592 | -9.6879903 |
| SSH-FN SSR-RM08 | Second Degree | 2.08024639 | -11.520954 |
| SSH-CH SSH-LE | Second Degree | 2.76801074 | -8.5077905 |
| SSH-LE SSH-NM | Second Degree | 4.14614921 | -7.2449034 |
| SSH-TY SSR-RM08 | Second Degree | 4.66020014 | -8.7724937 |
| SSH-FN SSH-TY | Second Degree | 7.72978808 | -5.8568765 |
| SSH-CH SSH-FG | Second Degree - BORDERLINE | NA | -0.9350357 |
| SSH-FN SSH-LU | Second Degree - BORDERLINE | NA | -0.9540334 |
| SSH-FN SSR-T5-1 | Second Degree - BORDERLINE | NA | -0.7513921 |
| SSH-KI SSH-SB | Second Degree - BORDERLINE | NA | -0.1614756 |
| SSR-RM08 SSR-T5-1 | Second Degree - BORDERLINE | NA | -1.184221 |
| SSH-TY SSR-T5-1 | Second Degree - BORDERLINE | NA | -0.6277514 |
| SSH-CH SSR-CR | Unrelated | NA | -18.5825 |
| SSH-CH SSH-FL | Unrelated | NA | -6.7345204 |
| SSH-CH SSH-FN | Unrelated | NA | -11.643368 |
| SSH-CH CNS-HE | Unrelated | NA | -23.334368 |
| SSH-CH SSH-KI | Unrelated | NA | -9.5097775 |
| SSH-CH SSH-LU | Unrelated | NA | -15.752172 |

| | | | |
|---|---|---|---|
| SSH-CH SSR-ML | Unrelated | NA | -21.216294 |
| SSH-CH SSH-NM | Unrelated | NA | -8.1536874 |
| SSH-CH SSH-PR | Unrelated | NA | -2.8890334 |
| SSH-CH SSH-RF | Unrelated | NA | -8.0697784 |
| SSH-CH SSR-RM08 | Unrelated | NA | -14.228357 |
| SSH-CH SSH-SB | Unrelated | NA | -6.9733214 |
| SSH-CH SSR-SN | Unrelated | NA | -9.5097705 |
| SSH-CH SSR-T5-1 | Unrelated | NA | -12.745411 |
| SSH-CH SSR-TH | Unrelated | NA | -10.951344 |
| SSH-CH SSH-TY | Unrelated | NA | -10.990837 |
| SSR-CRCNS-HE | Unrelated | NA | -30.384998 |
| SSR-CRSSR-RM08 | Unrelated | NA | -16.865005 |
| SSR-CRSSR-T5-1 | Unrelated | NA | -17.151901 |
| SSR-CRSSR-TH | Unrelated | NA | -18.925005 |
| SSH-FL SSR-CR | Unrelated | NA | -18.975925 |
| SSH-FL SSH-FN | Unrelated | NA | -13.270147 |
| SSH-FL CNS-HE | Unrelated | NA | -25.702595 |
| SSH-FL SSH-KI | Unrelated | NA | -8.7067172 |
| SSH-FL SSH-LE | Unrelated | NA | -12.793677 |
| SSH-FL SSH-LU | Unrelated | NA | -16.082093 |
| SSH-FL SSR-ML | Unrelated | NA | -20.823515 |
| SSH-FL SSH-NM | Unrelated | NA | -15.307407 |
| SSH-FL SSH-PR | Unrelated | NA | -11.279536 |
| SSH-FL SSR-RM08 | Unrelated | NA | -14.931369 |
| SSH-FL SSH-SB | Unrelated | NA | -7.3355896 |
| SSH-FL SSR-SN | Unrelated | NA | -12.965839 |
| SSH-FL SSR-T5-1 | Unrelated | NA | -14.10311 |

| | | | |
|---|---|---|---|
| SSH-FL SSR-TH | Unrelated | NA | -14.461288 |
| SSH-FL SSH-TY | Unrelated | NA | -11.095807 |
| SSH-FG SSR-CR | Unrelated | NA | -16.159919 |
| SSH-FG SSH-FN | Unrelated | NA | -11.81085 |
| SSH-FG CNS-HE | Unrelated | NA | -28.332142 |
| SSH-FG SSH-KI | Unrelated | NA | -6.0262484 |
| SSH-FG SSH-LU | Unrelated | NA | -14.703535 |
| SSH-FG SSR-ML | Unrelated | NA | -17.547706 |
| SSH-FG SSH-NM | Unrelated | NA | -2.8287297 |
| SSH-FG SSH-RF | Unrelated | NA | -14.278475 |
| SSH-FG SSR-RM08 | Unrelated | NA | -14.083598 |
| SSH-FG SSH-SB | Unrelated | NA | -5.5300827 |
| SSH-FG SSR-SN | Unrelated | NA | -12.784172 |
| SSH-FG SSR-T5-1 | Unrelated | NA | -11.433419 |
| SSH-FG SSR-TH | Unrelated | NA | -13.424443 |
| SSH-FG SSH-TY | Unrelated | NA | -11.203803 |
| SSH-FN SSR-CR | Unrelated | NA | -14.70078 |
| SSH-FN CNS-HE | Unrelated | NA | -29.896915 |
| SSH-FN SSH-KI | Unrelated | NA | -12.316069 |
| SSH-FN SSH-LE | Unrelated | NA | -14.568492 |
| SSH-FN SSR-ML | Unrelated | NA | -15.357709 |
| SSH-FN SSH-NM | Unrelated | NA | -12.156449 |
| SSH-FN SSH-PR | Unrelated | NA | -10.880441 |
| SSH-FN SSH-RF | Unrelated | NA | -17.012475 |
| SSH-FN SSH-SB | Unrelated | NA | -8.5681839 |
| SSH-FN SSR-SN | Unrelated | NA | -11.322524 |
| SSH-FN SSR-TH | Unrelated | NA | -19.483514 |

| | | | |
|---|---|---|---|
| SSH-KI SSR-CR | Unrelated | NA | -14.443782 |
| SSH-KI CNS-HE | Unrelated | NA | -25.762135 |
| SSH-KI SSH-LE | Unrelated | NA | -10.286858 |
| SSH-KI SSH-LU | Unrelated | NA | -13.721384 |
| SSH-KI SSR-ML | Unrelated | NA | -16.356312 |
| SSH-KI SSH-NM | Unrelated | NA | -10.749899 |
| SSH-KI SSH-PR | Unrelated | NA | -3.9170118 |
| SSH-KI SSH-RF | Unrelated | NA | -12.508703 |
| SSH-KI SSR-RM08 | Unrelated | NA | -13.290003 |
| SSH-KI SSR-SN | Unrelated | NA | -9.7590083 |
| SSH-KI SSR-T5-1 | Unrelated | NA | -11.247495 |
| SSH-KI SSR-TH | Unrelated | NA | -17.080913 |
| SSH-KI SSH-TY | Unrelated | NA | -5.3915938 |
| SSH-LE SSR-CR | Unrelated | NA | -15.257042 |
| SSH-LE CNS-HE | Unrelated | NA | -24.918661 |
| SSH-LE SSH-LU | Unrelated | NA | -13.946423 |
| SSH-LE SSR-ML | Unrelated | NA | -16.197402 |
| SSH-LE SSH-RF | Unrelated | NA | -9.7073574 |
| SSH-LE SSR-RM08 | Unrelated | NA | -16.564771 |
| SSH-LE SSH-SB | Unrelated | NA | -13.809711 |
| SSH-LE SSR-SN | Unrelated | NA | -9.1798704 |
| SSH-LE SSR-T5-1 | Unrelated | NA | -14.139968 |
| SSH-LE SSR-TH | Unrelated | NA | -8.0925669 |
| SSH-LE SSH-TY | Unrelated | NA | -11.320277 |
| SSH-LU SSR-CR | Unrelated | NA | -14.192507 |
| SSH-LU CNS-HE | Unrelated | NA | -27.834796 |
| SSH-LU SSR-ML | Unrelated | NA | -13.451977 |

| | | | |
|---|---|---|---|
| SSH-LU SSH-NM | Unrelated | NA | -11.200368 |
| SSH-LU SSH-PR | Unrelated | NA | -11.687948 |
| SSH-LU SSH-RF | Unrelated | NA | -17.297913 |
| SSH-LU SSR-RM08 | Unrelated | NA | -13.775198 |
| SSH-LU SSH-SB | Unrelated | NA | -12.818399 |
| SSH-LU SSR-SN | Unrelated | NA | -10.908026 |
| SSH-LU SSR-T5-1 | Unrelated | NA | -8.9450829 |
| SSH-LU SSR-TH | Unrelated | NA | -18.108024 |
| SSH-LU SSH-TY | Unrelated | NA | -11.38017 |
| SSR-MLCNS-HE | Unrelated | NA | -34.015276 |
| SSR-MLSSR-RM08 | Unrelated | NA | -19.68116 |
| SSR-MLSSR-T5-1 | Unrelated | NA | -19.096509 |
| SSR-MLSSR-TH | Unrelated | NA | -21.462774 |
| SSH-NM SSR-CR | Unrelated | NA | -6.4282148 |
| SSH-NM CNS-HE | Unrelated | NA | -26.202074 |
| SSH-NM SSR-ML | Unrelated | NA | -10.704904 |
| SSH-NM SSH-PR | Unrelated | NA | -3.5299099 |
| SSH-NM SSH-RF | Unrelated | NA | -15.372697 |
| SSH-NM SSR-RM08 | Unrelated | NA | -15.223125 |
| SSH-NM SSH-SB | Unrelated | NA | -11.680437 |
| SSH-NM SSR-SN | Unrelated | NA | -10.478441 |
| SSH-NM SSR-T5-1 | Unrelated | NA | -15.303962 |
| SSH-NM SSR-TH | Unrelated | NA | -7.2518247 |
| SSH-NM SSH-TY | Unrelated | NA | -11.395118 |
| SSH-PR SSR-CR | Unrelated | NA | -13.492014 |
| SSH-PR CNS-HE | Unrelated | NA | -22.741725 |
| SSH-PR SSR-ML | Unrelated | NA | -13.627337 |

| | | | |
|---|---|---|---|
| SSH-PR SSH-RF | Unrelated | NA | -10.530447 |
| SSH-PR SSR-RM08 | Unrelated | NA | -11.925433 |
| SSH-PR SSR-SN | Unrelated | NA | -7.0473026 |
| SSH-PR SSR-T5-1 | Unrelated | NA | -11.336723 |
| SSH-PR SSR-TH | Unrelated | NA | -11.827428 |
| SSH-PR SSH-TY | Unrelated | NA | -8.1238338 |
| SSH-RF SSR-CR | Unrelated | NA | -19.820108 |
| SSH-RF CNS-HE | Unrelated | NA | -21.927939 |
| SSH-RF SSR-ML | Unrelated | NA | -22.158346 |
| SSH-RF SSR-RM08 | Unrelated | NA | -20.428123 |
| SSH-RF SSH-SB | Unrelated | NA | -14.913188 |
| SSH-RF SSR-SN | Unrelated | NA | -11.575057 |
| SSH-RF SSR-T5-1 | Unrelated | NA | -16.356174 |
| SSH-RF SSR-TH | Unrelated | NA | -13.526404 |
| SSH-RF SSH-TY | Unrelated | NA | -13.669551 |
| SSR-RM08CNS-HE | Unrelated | NA | -33.004656 |
| SSR-RM08SSR-TH | Unrelated | NA | -21.052304 |
| SSH-SB SSR-CR | Unrelated | NA | -13.803784 |
| SSH-SB CNS-HE | Unrelated | NA | -25.964633 |
| SSH-SB SSR-ML | Unrelated | NA | -16.782508 |
| SSH-SB SSR-RM08 | Unrelated | NA | -11.824206 |
| SSH-SB SSR-SN | Unrelated | NA | -8.7107269 |
| SSH-SB SSR-T5-1 | Unrelated | NA | -11.766951 |
| SSH-SB SSR-TH | Unrelated | NA | -17.758238 |
| SSH-SB SSH-TY | Unrelated | NA | -9.0353101 |
| SSR-SN SSR-CR | Unrelated | NA | -15.396173 |
| SSR-SN CNS-HE | Unrelated | NA | -19.884924 |

| | | | |
|---|---|---|---|
| SSR-SN SSR-ML | Unrelated | NA | -14.615784 |
| SSR-SN SSR-RM08 | Unrelated | NA | -13.609378 |
| SSR-SN SSR-T5-1 | Unrelated | NA | -12.435657 |
| SSR-SN SSR-TH | Unrelated | NA | -15.025578 |
| SSR-SN SSH-TY | Unrelated | NA | -10.864574 |
| SSR-T5-1CNS-HE | Unrelated | NA | -29.122807 |
| SSR-T5-1SSR-TH | Unrelated | NA | -17.969181 |
| SSR-THCNS-HE | Unrelated | NA | -29.104437 |
| SSH-TY SSR-CR | Unrelated | NA | -12.695972 |
| SSH-TY CNS-HE | Unrelated | NA | -28.065183 |
| SSH-TY SSR-ML | Unrelated | NA | -14.636868 |
| SSH-TY SSR-TH | Unrelated | NA | -16.291348 |

### SUPPLEMENTAL FIGURES

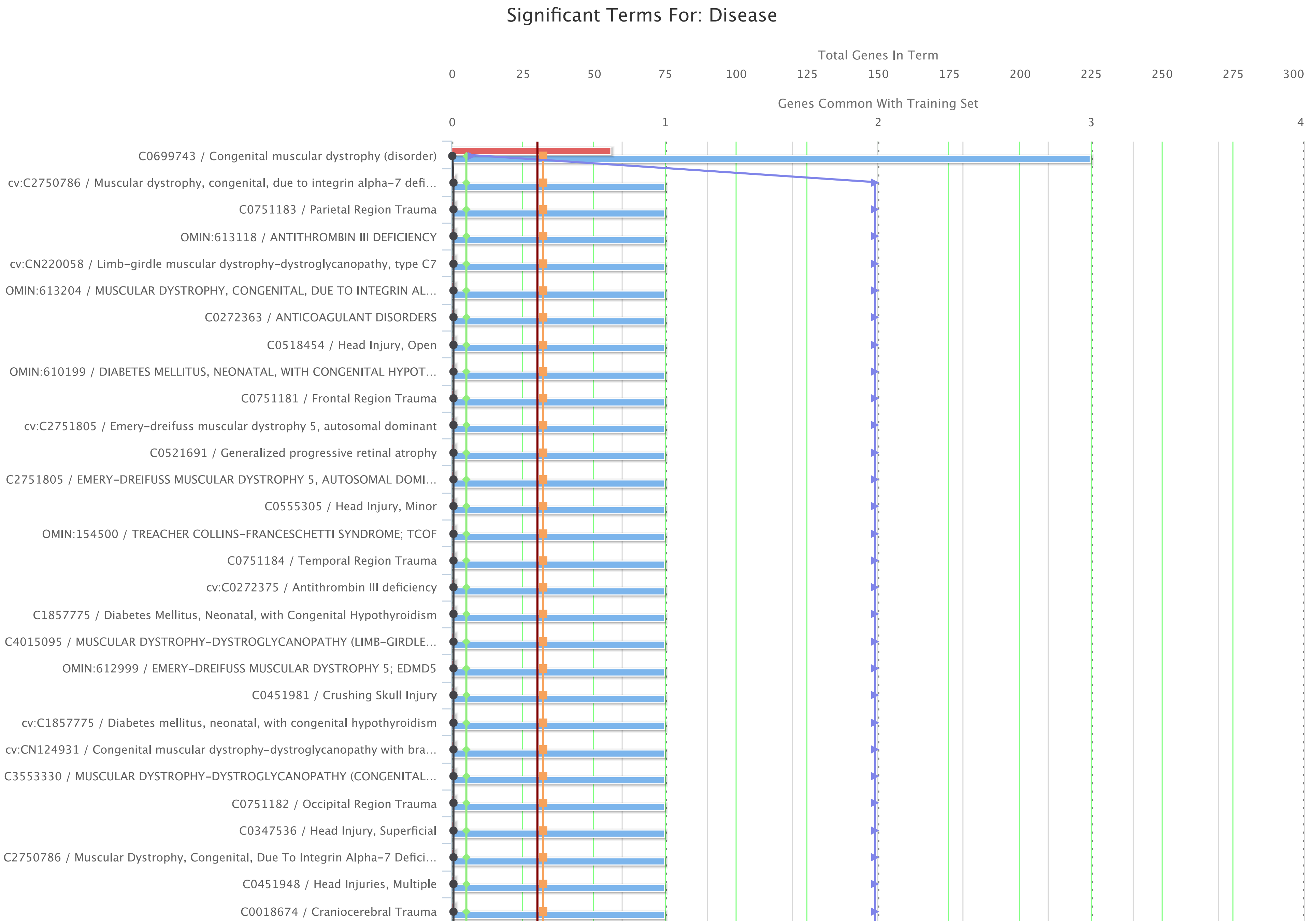

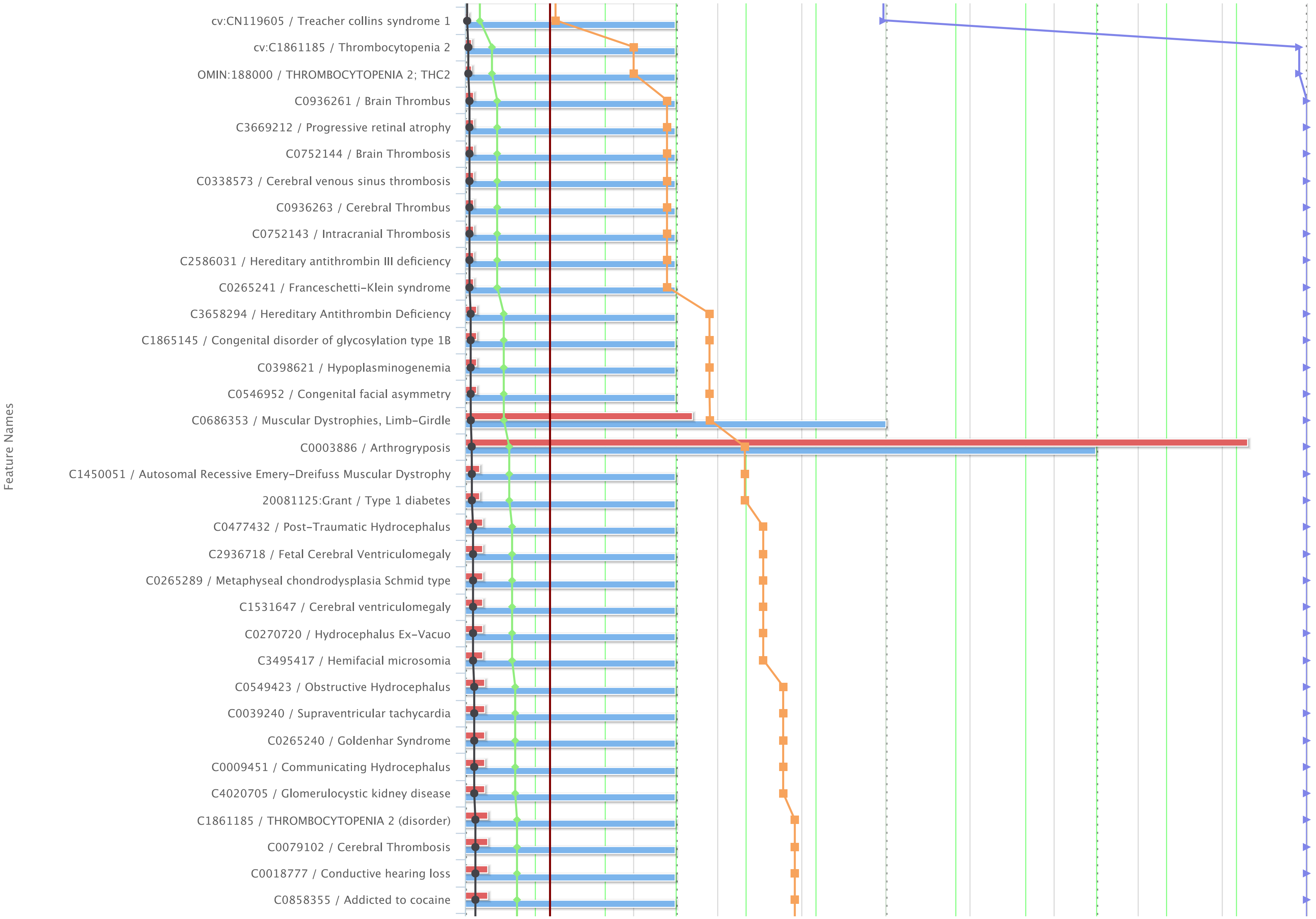
Feature Names
cv:CN119605 / Treacher collins syndrome 1
cv:C1861185 / Thrombocytopenia 2
OMIN:188000 / THROMBOCYTOPENIA 2; THC2
C0936261 / Brain Thrombus
C3669212 / Progressive retinal atrophy
C0752144 / Brain Thrombosis
C0338573 / Cerebral venous sinus thrombosis
C0936263 / Cerebral Thrombus
C0752143 / Intracranial Thrombosis
C2586031 / Hereditary antithrombin III deficiency
C0265241 / Franceschetti-Klein syndrome
C3658294 / Hereditary Antithrombin Deficiency
C1865145 / Congenital disorder of glycosylation type 1B
C0398621 / Hypoplasminogenemia
C0546952 / Congenital facial asymmetry
C0686353 / Muscular Dystrophies, Limb-Girdle
C0003886 / Arthrogryposis
C1450051 / Autosomal Recessive Emery-Dreifuss Muscular Dystrophy
20081125:Grant / Type 1 diabetes
C0477432 / Post-Traumatic Hydrocephalus
C2936718 / Fetal Cerebral Ventriculomegaly
C0265289 / Metaphyseal chondrodysplasia Schmid type
C1531647 / Cerebral ventriculomegaly
C0270720 / Hydrocephalus Ex-Vacuo
C3495417 / Hemifacial microsomia
C0549423 / Obstructive Hydrocephalus
C0039240 / Supraventricular tachycardia
C0265240 / Goldenhar Syndrome
C0009451 / Communicating Hydrocephalus
C4020705 / Glomerulocystic kidney disease
C1861185 / THROMBOCYTOPENIA 2 (disorder)
C0079102 / Cerebral Thrombosis
C0018777 / Conductive hearing loss
C0858355 / Addicted to cocaine

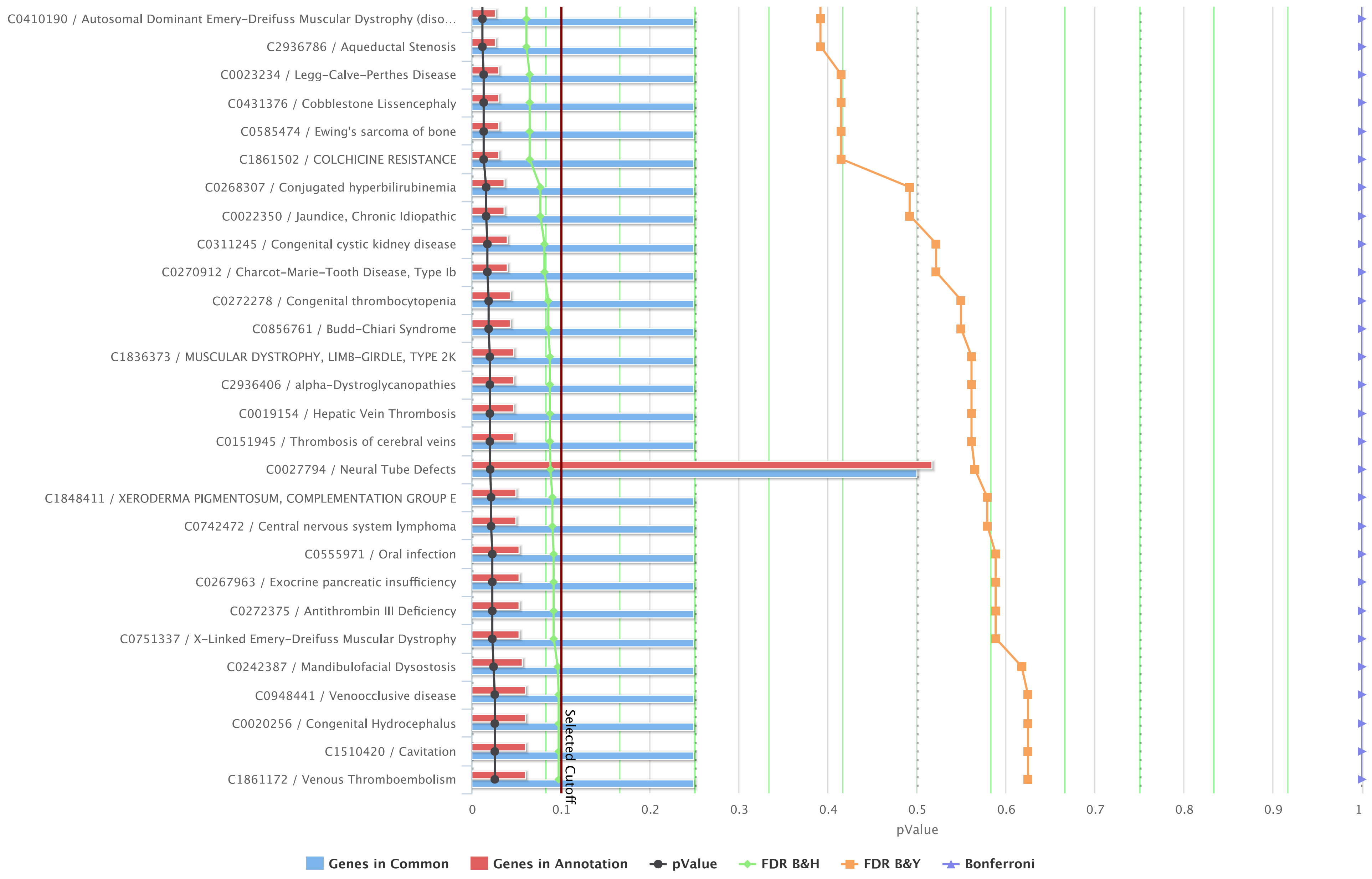

**Figure S1:** Significant terms for disease identified by ToppFun scan of genes within high FST (top 0.5%).

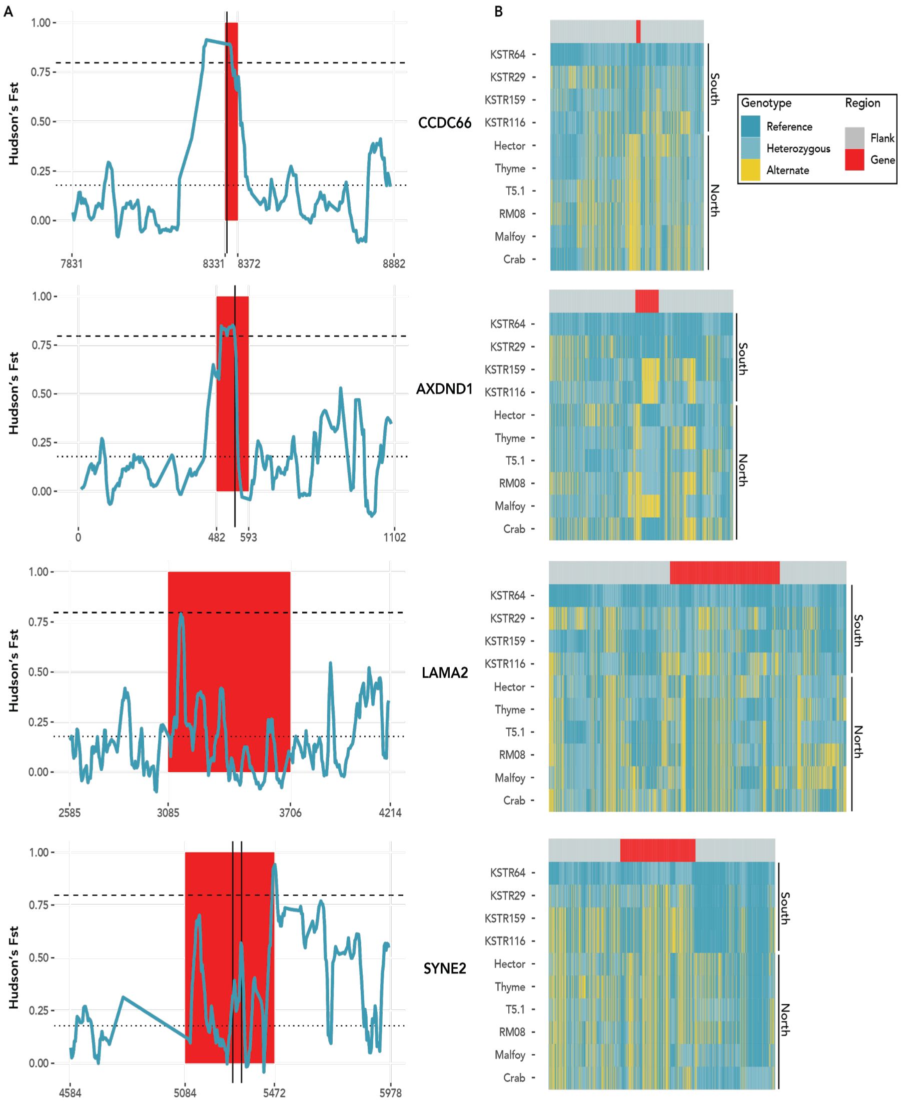

**Figure S2**: Additional highly differentiated genes between wet and dry forest populations referenced in the text. A: Hudson's $F_{ST}$ within windows of 20kb with a 4 kb slide. Gene regions are in red, flanked by 500 kb (or length to beginning or end of scaffold) of sequence. X-axis values correspond to position along the scaffold. The dotted line indicates average $F_{ST}$ value across all windows ($F_{ST}$ = 0.178), and the dashed line represents the top 0.5% of values ($F_{ST}$ = 0.797). Vertical black lines indicate a non-synonymous SNP with an Fst >= 0.750, excluding BCAS3 (see Results). B: Heatmaps indicating the pattern of SNP variation within and surrounding highly divergent genes. SNVs within the genes are located under the red band and those within 200 kb of flanking region under the gray bands.

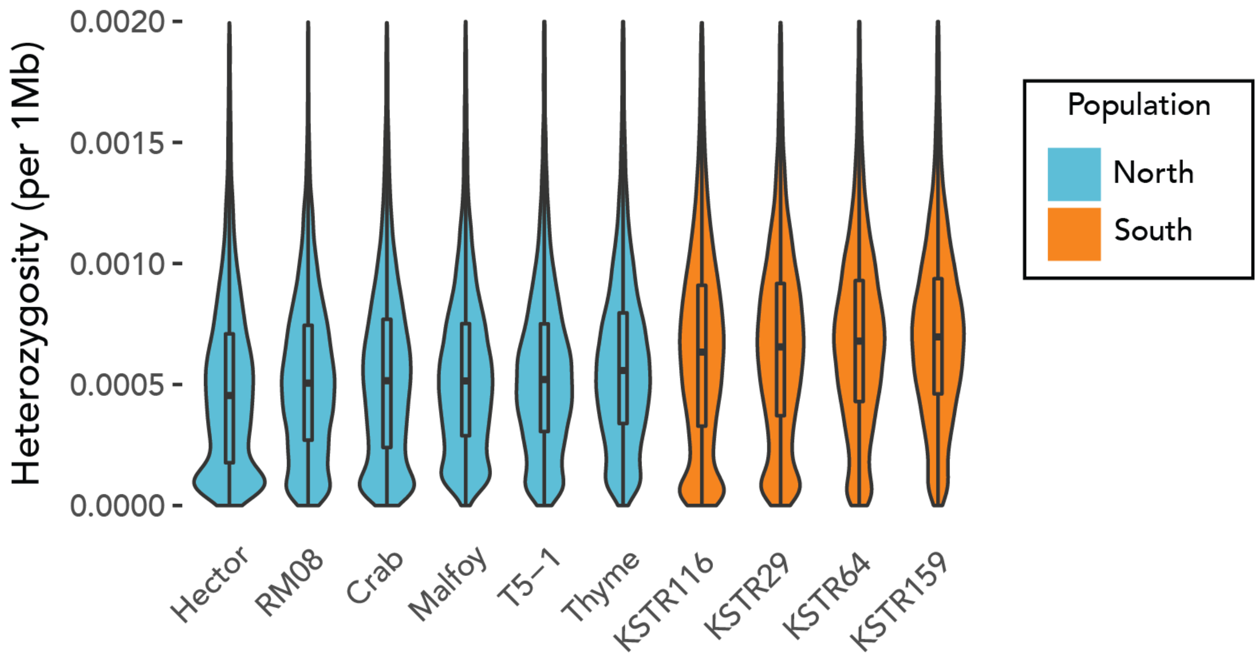

**Figure S3**: Violin and box plots of heterozygosity values in 1Mb / 200 Kb sliding windows sorted from lowest to highest median value for the 10 high-coverage samples. The individuals from the southern population consistently have higher values (W = 1,535,400,000, p < 2.2e-16).

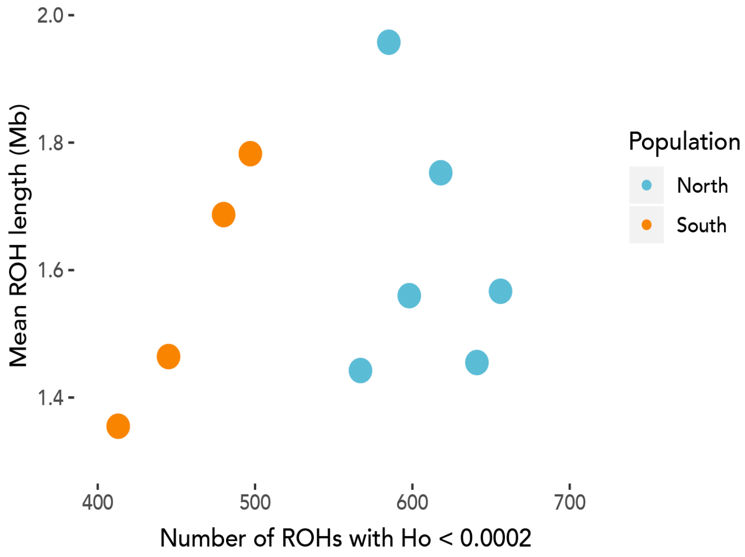

**Figure S4**: Mean length and total number of runs of homozygosity (≥ 1 Mb with Ho < 0.0002) per individual in 1 Mb / 200 Kb sliding windows. Although the mean length of ROHs overlaps between populations, there are significantly more ROHs in the northern population (W = 24, p-value = 0.009524)

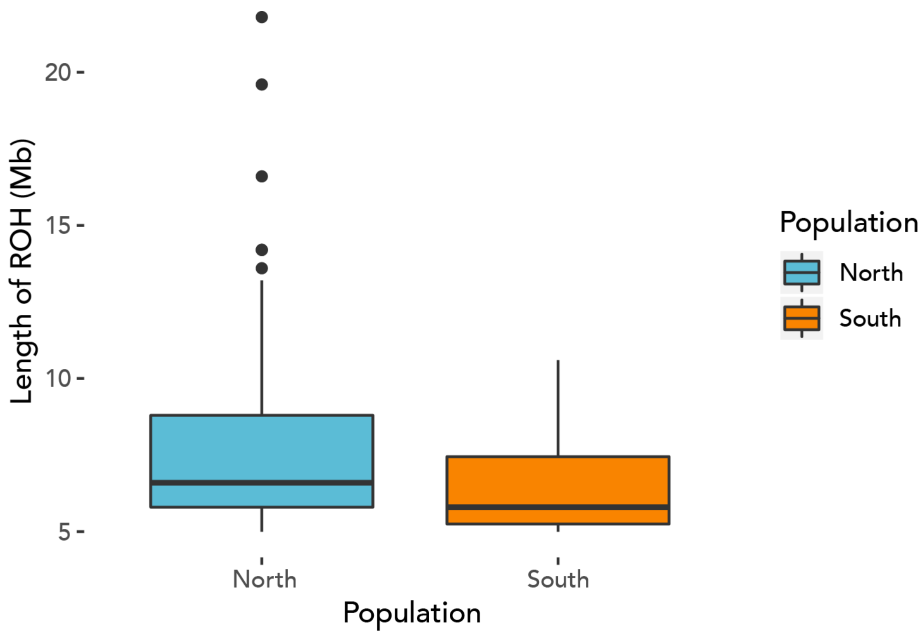

**Figure S5**: Difference in length of extended long runs of homozygosity (≥ 1 Mb with Ho < 0.0002 for at least 5 Mb); (W = 1315.5, p-value = 0.0243).

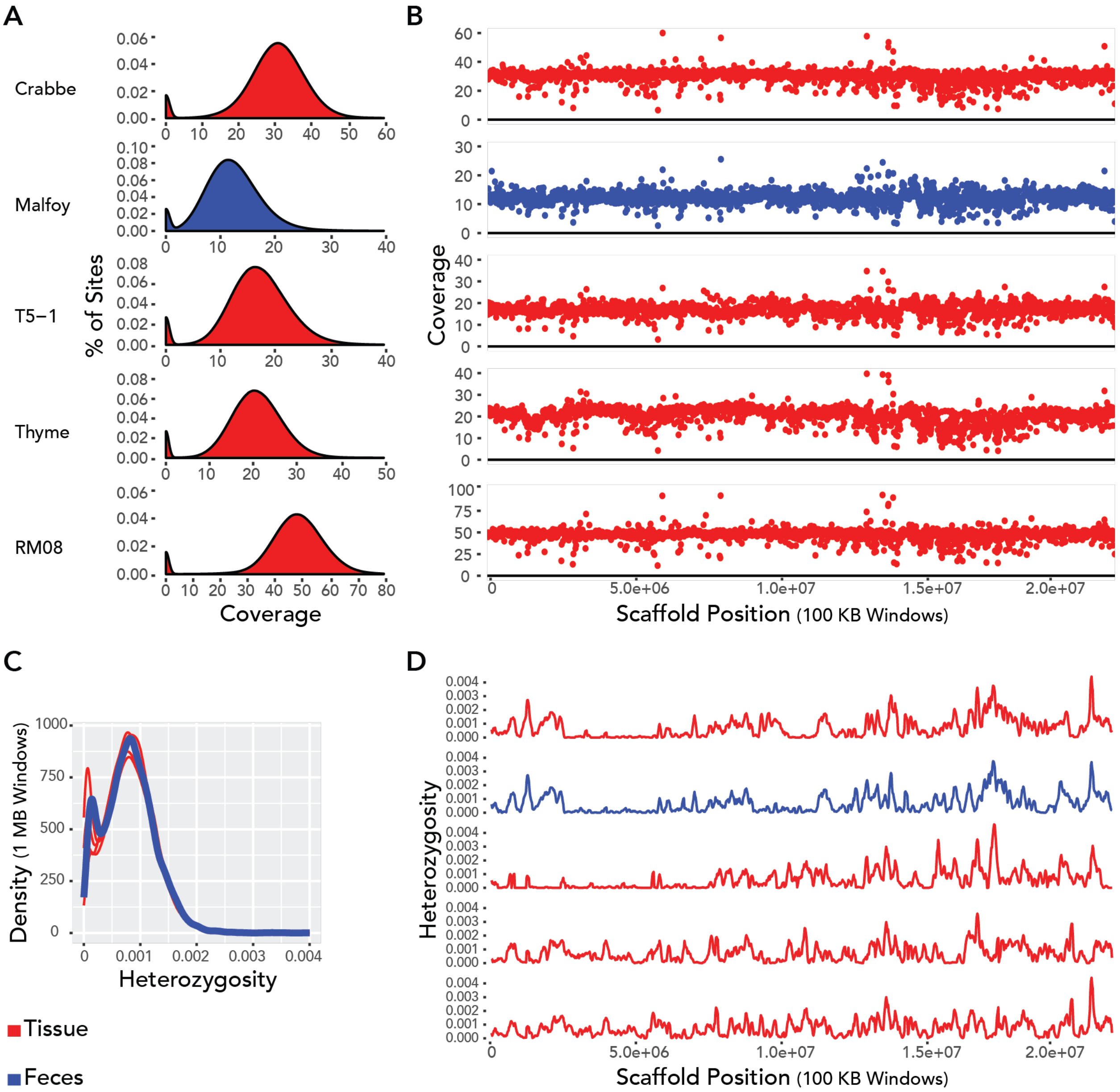

**Figure S6:** A) Density of genomic coverage of high-coverage genomes from Santa Rosa. B) Average coverage per 100 KB window along the largest scaffold of the *C. imitator* 1.0 reference genome. C) Density of 1 MB windows at varying levels of heterozygosity along the entire genome. D) Heterozygosity of 100 KB windows along the largest scaffold of the *C. imitator* 1.0 reference genome. The top two genomes (SSR-CR and SSR-ML) are from siblings. The order of individuals in figures B and D corresponds to that of figure A.

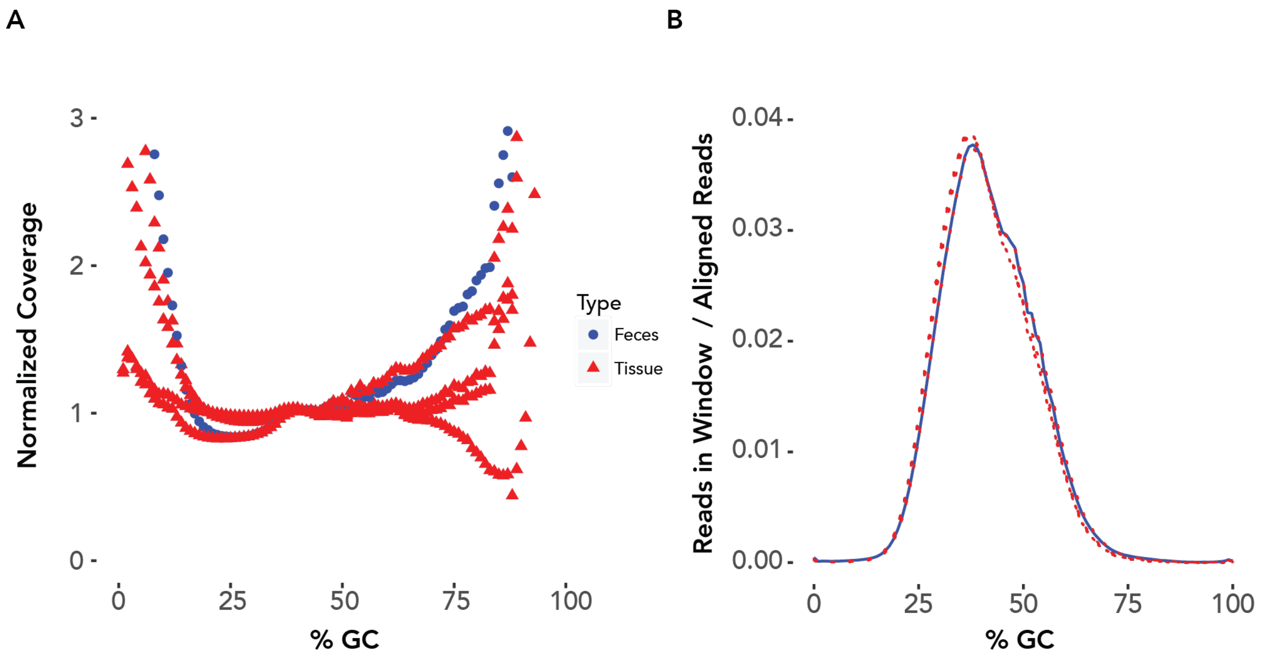

**Figure S7:** Percent of GC content across the genome for the four tissue (red) and one fecal (blue) samples from Sector Santa Rosa. GC content does not substantially differ for each type of sample. A) Average normalized coverage at each percentage of GC. B) Number of reads per 100 bp window (scaled by the number aligned reads) at each percentage of GC.

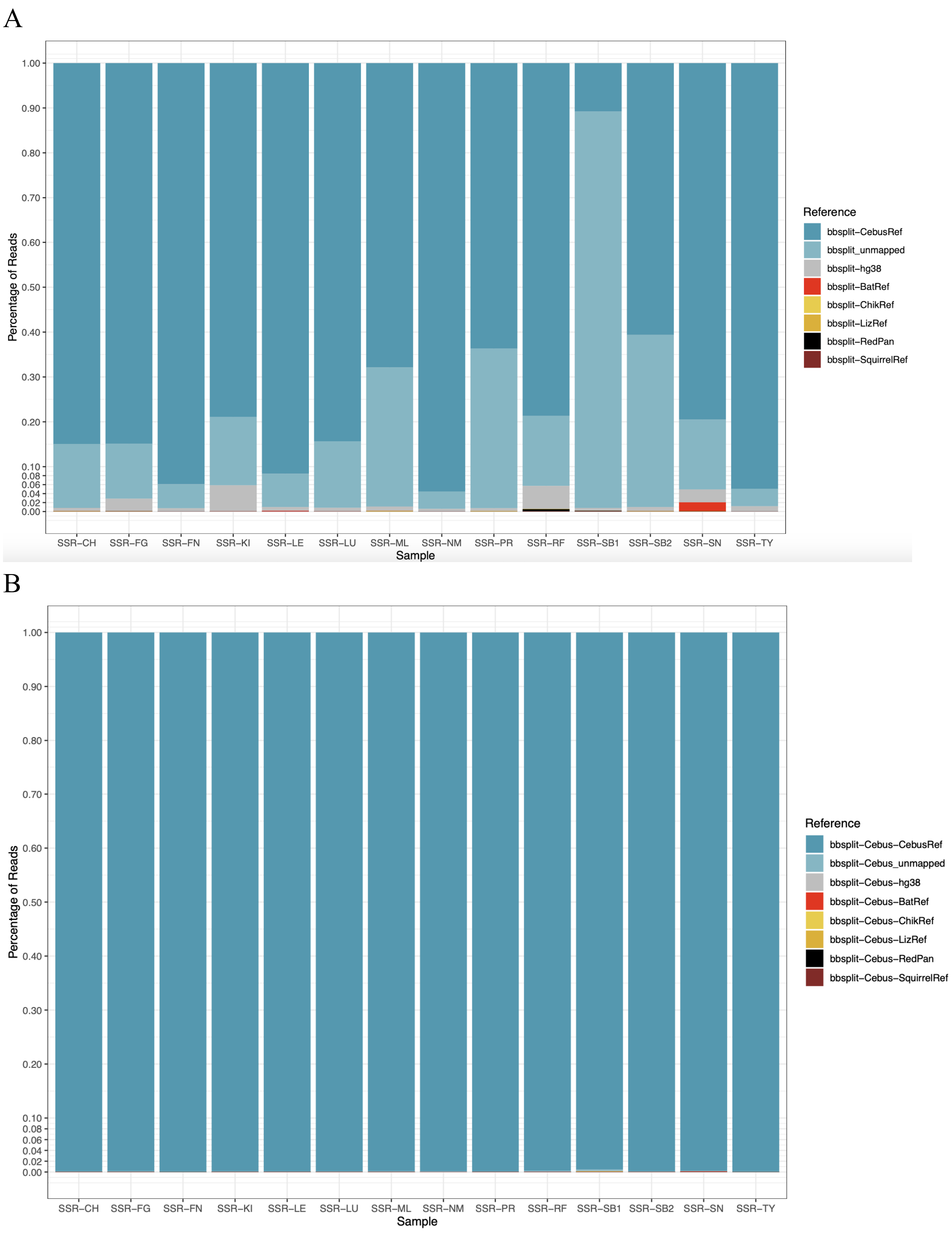

**Figure S8:** A: The percentages of fecalFACS reads mapping to *Cebus imitator*, hg36, and 5 genomes, used to estimate the contribution of prey items (bat, bird, lizard, coati, squirrel): *Phyllostomus discolor*, *Gallus gallus*, *Pogona vitticeps*, *Ailurus fulgens*, *Sciurus carolinensis*, respectively. B: The percentage of reads mapping to each of the 7 genomes after bbsplit was used to select only the *Cebus imitator* reads, which demonstrates that effectively no contaminating or dietary reads entered the fecalFACS analysis

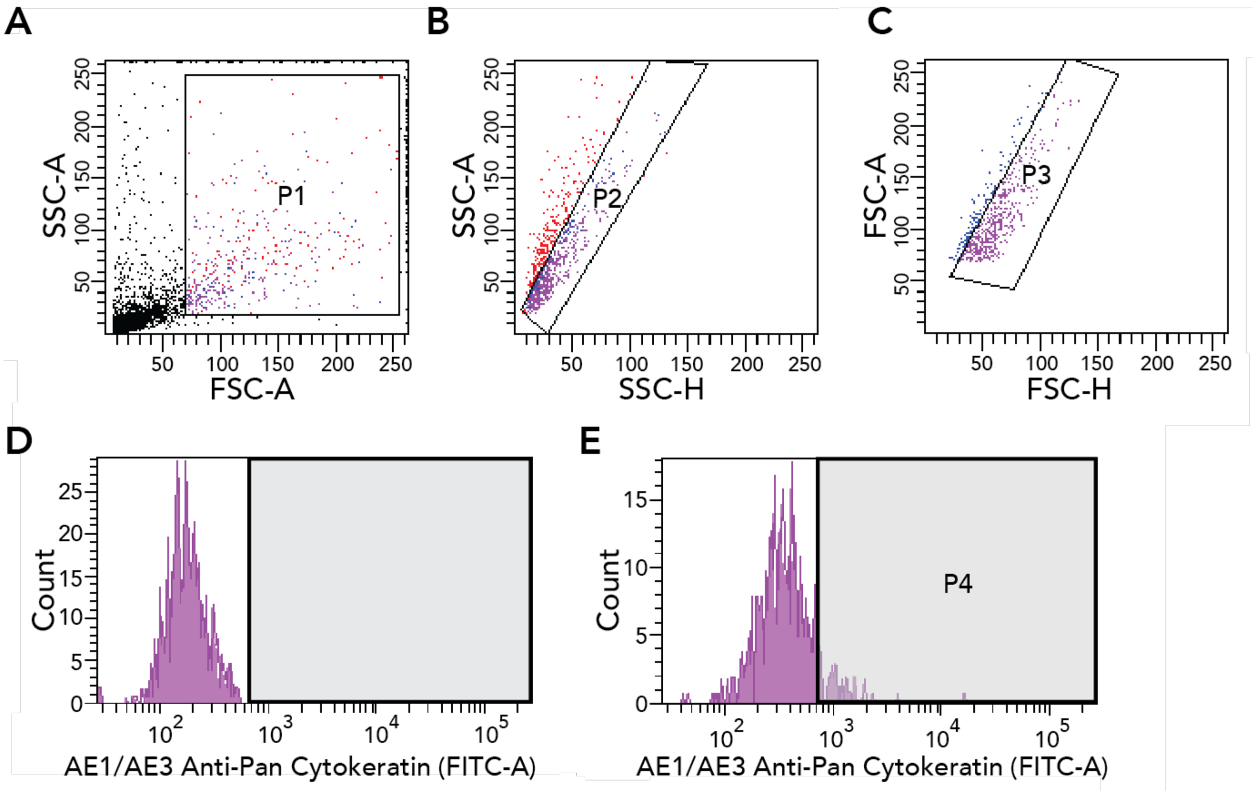

**Figure S9:** FACS gating strategy. Cells were gated first by size and complexity to avoid bacteria and cellular debris (A), followed by discrimination of cellular agglomerations (B and C). Fluorescence of AE1/AE3 Pan Cytokeratin Alexa Fluor® 488 antibody (FITC-A) is depicted in unstained (D) and stained (E) cellular populations. Epithelial cells were identified as those fluorescing beyond background levels, as depicted in the P4 gate. Note: images have been altered in a slight but non-material manner from raw output to improve labeling and visibility of gating strategy for schematic purposes.

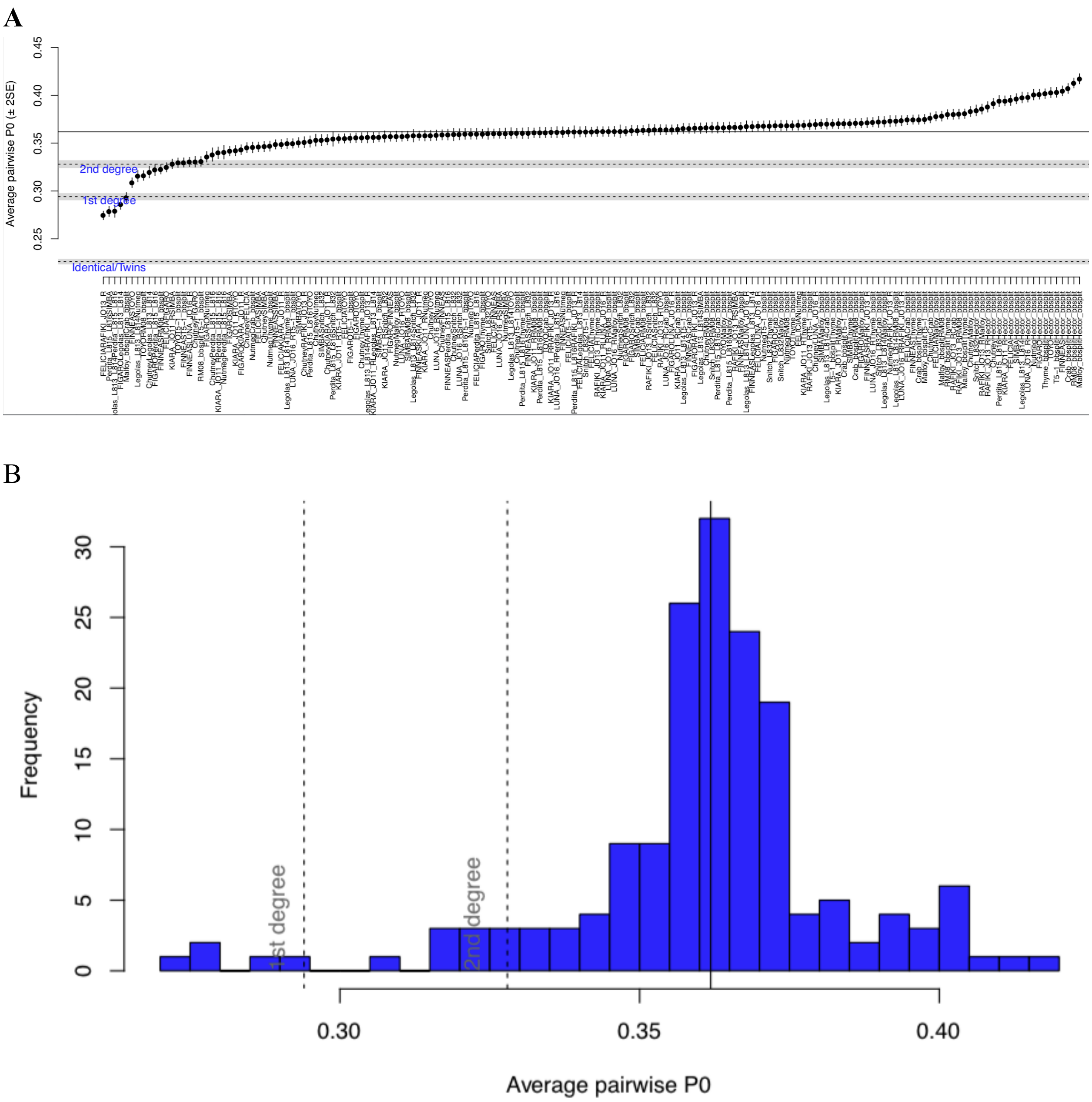

**Figure S10:** A: Pairwise estimates of relatedness generated from READ (21). B: Distribution of relatedness scores

#### REFERENCES

1. S. Gnerre, *et al.*, High-quality draft assemblies of mammalian genomes from massively parallel sequence data. *Proc. Natl. Acad. Sci. U. S. A.* **108**, 1513–1518 (2011).
2. A. Morgulis, E. M. Gertz, A. A. Schäffer, R. Agarwala, WindowMasker: window-based masker for sequenced genomes. *Bioinformatics* **22**, 134–141 (2006).
3. E. S. Rice, R. E. Green, New Approaches for Genome Assembly and Scaffolding. *Annu Rev Anim Biosci* **7**, 17–40 (2019).
4. A. M. Bolger, M. Lohse, B. Usadel, Trimmomatic: a flexible trimmer for Illumina sequence data. *Bioinformatics* **30**, 2114–2120 (2014).
5. H. Li, R. Durbin, Fast and accurate short read alignment with Burrows-Wheeler transform. *Bioinformatics* **25**, 1754–1760 (2009).
6. H. Li, *et al.*, The Sequence Alignment/Map format and SAMtools. *Bioinformatics* **25**, 2078–2079 (2009).
7. A. McKenna, *et al.*, The Genome Analysis Toolkit: a MapReduce framework for analyzing next-generation DNA sequencing data. *Genome Res.* **20**, 1297–1303 (2010).
8. L. Fu, B. Niu, Z. Zhu, S. Wu, W. Li, CD-HIT: accelerated for clustering the next-generation sequencing data. *Bioinformatics* **28**, 3150–3152 (2012).
9. A. M. Altenhoff, *et al.*, OMA standalone: orthology inference among public and custom genomes and transcriptomes. *Genome Res.* **29**, 1152–1163 (2019).
10. S. Kumar, G. Stecher, M. Suleski, S. B. Hedges, TimeTree: A resource for timelines, timetrees, and divergence times. *Mol. Biol. Evol.* **34**, 1812–1819 (2017).
11. J. W. Brown, J. F. Walker, S. A. Smith, Phyx: phylogenetic tools for unix. *Bioinformatics* **33**, 1886–1888 (2017).
12. K. Katoh, D. M. Standley, MAFFT multiple sequence alignment software version 7: improvements in performance and usability. *Mol. Biol. Evol.* **30**, 772–780 (2013).
13. M. J. Hubisz, K. S. Pollard, A. Siepel, PHAST and RPHAST: phylogenetic analysis with space/time models. *Brief. Bioinform.* **12**, 41–51 (2011).
14. M. Anisimova, J. P. Bielawski, Z. Yang, Accuracy and power of the likelihood ratio test in detecting adaptive molecular evolution. *Mol. Biol. Evol.* **18**, 1585–1592 (2001).
15. Z. Yang, PAML 4: phylogenetic analysis by maximum likelihood. *Mol. Biol. Evol.* **24**, 1586–1591 (2007).
16. J. Huerta-Cepas, F. Serra, P. Bork, ETE 3: Reconstruction, Analysis, and Visualization of

Phylogenomic Data. *Mol. Biol. Evol.* **33**, 1635–1638 (2016).

17. J. Zhang, Evaluation of an Improved Branch-Site Likelihood Method for Detecting Positive Selection at the Molecular Level. *Molecular Biology and Evolution* **22**, 2472–2479 (2005).

18. J. Chen, E. E. Bardes, B. J. Aronow, A. G. Jegga, ToppGene Suite for gene list enrichment analysis and candidate gene prioritization. *Nucleic Acids Res.* **37**, W305–11 (2009).

19. R. Tacutu, *et al.*, Human Ageing Genomic Resources: new and updated databases. *Nucleic Acids Res.* **46**, D1083–D1090 (2018).

20. S. Hayden, *et al.*, A cluster of olfactory receptor genes linked to frugivory in bats. *Mol. Biol. Evol.* **31**, 917–927 (2014).

21. J. M. Monroy Kuhn, M. Jakobsson, T. Günther, Estimating genetic kin relationships in prehistoric populations. *PLoS One* **13**, e0195491 (2018).
